# Characterization of membrane structures regulating primary ciliogenesis by quantitative isotropic ultrastructure imaging

**DOI:** 10.1101/2025.08.20.670930

**Authors:** Quanlong Lu, Huijie Zhao, Ziam Khan, Adam Harned, Erina Kamiya, Valentin Magidson, Abhi Senthilkumar, Sumeth Perera, Kedar Narayan, Christopher J. Westlake

## Abstract

The trafficking, docking, and fusion of membrane vesicles at the mother centriole (MC) are required to construct the primary cilium. Here, we determined the three-dimensional (3D) membrane ultrastructures, and associated proteins, involved in primary cilium assembly upstream of axoneme growth. Our work reveals that the enlargement of small vesicles docked to the MC is a key trigger for ciliogenesis progression, a process requiring the MC distal appendage protein CEP164. We show these vesicles subsequently fuse to form tubular C-shaped and an unprecedented toroidal membrane intermediates, which ultimately organize into the ciliary vesicle covering the MC distal end. The formation of these previously uncharacterized tubular membrane ciliogenesis intermediates is orchestrated by the membrane trafficking regulators EHD1 and RAB8, and requires the IFT-B complex protein IFT88. Remarkably, we show that EHD1, through its membrane tubulation function, regulates ciliogenesis progression by directly promoting CP110/CEP97 removal from the MC cap. The establishment of these tubular membrane structures is also associated with the recruitment of the ciliary gate transition zone proteins. This study changes the architectural framework for understanding ciliogenesis mechanisms and highlights the application of isotropic ultrastructure imaging and three-dimensional quantitative analysis in understanding membrane trafficking and organelle biogenesis mechanisms.

## Introduction

Primary cilia extend from the surface of most eukaryotic cells, serving as sensory structures important for detecting and transducing mechanical and chemical signals which are essential for embryonic development, cellular polarity, and organogenesis^1,2,3^. Impairment in the formation and function of primary cilia causes human disease that can affect multiple organs and tissues^1,2,3^. The primary cilium develops from the mother centriole (MC) and consists of a microtubule-based axoneme surrounded by a ciliary membrane^3^. The process of ciliogenesis is a tightly regulated sequence of events involving MC to basal body (BB) transition, transition zone (TZ) formation, and axoneme assembly^2,4^. The initial stages of cilium biogenesis occur in the cytoplasm and requires the transport of preciliary vesicles (PCV) to the MC^4,5,6,7^. These membranes dock to the distal appendages (DA) on the MC and are referred to as DA vesicles (DAV). The DA proteins (DAP) are crucial for cilia formation and have been proposed to mediate the docking of DAVs to the MC by interacting with proteins on PCVs^8,9,10^. The hallmark of early ciliogenesis is a ∼300 nm ciliary vesicle (CV)^4,11^ that covers the distal end of the MC that is assembled from smaller DAVs^5^. The intraflagellar transport (IFT)-B complex functions in axoneme formation at the CV stage^5^, and this vesicular structure reorganizes into a double membrane sheath around the extending axoneme. Initiation of axoneme growth requires the proteolytic removal of CP110 and CEP97, the MC cap, from the MC distal end via a process facilitated by DAPs^12,13,14^ and linked to the establishment of the CV ^2,5,15^. TZ protein recruitment to the MC is also associated with the CV stage^5,16^ and leads to establishment of the TZ at the base of the cilium which serves to regulate ciliary transport.

Several membrane trafficking regulators have been identified as crucial for primary ciliogenesis, including members of the RAB small GTPase family^4,17,18,19,20,21^. RAB8 was the first RAB shown to function in ciliogenesis and was linked to assembly from the CV stage^5,22,23^. A RAB cascade involving RAB11 and RAB8 is integral to this process. RAB11 facilitates the vesicular transport of RABIN8, a RAB8 guanine nucleotide exchange factor (GEF), to the MC resulting in RAB8 activation^6,24^. Highlighting a complex poorly understood mechanism of membrane trafficking regulation of ciliogenesis RAB34 is also essential for CV assembly^25,26,27,28^ and RAB19 has been linked to early stages of primary cilium assembly^29^. The RAB-associated membrane shaping and fusion proteins EH domain-containing (EHD) proteins EHD1 and EHD3, and their associated factors PACSIN1/2, SNAP29 and MICAL-L1 are also important for pre-CV stages of ciliogenesis^5,16,30^. Additionally, EHD1 and PACSIN1 membrane tubulation function is important for the fusion of the CV and ciliary sheath membranes with the plasma membrane (PM) via the formation of extracellular membrane channels (EMC), thus exposing the cilia membrane to the extracellular region^16^. Notably, EHD1, PACSIN1 and MICAL-L1 are essential for the removal of CP110/CEP97 from the MC cap^5,15,16,30,31^. How these trafficking regulators and associated membranes coordinate to initiate ciliogenesis on the MC through assembly of the CV is not clear.

Transmission electron microscopy (TEM) imaging has contributing greatly to our understanding of ciliogenesis pathways having been used to identify all the known early ciliogenesis intermediate structures, such as DAV, CV, and ciliary sheath^2,4,11,12,32^. However, TEM is limited to two-dimensional (2D) imaging of ∼70-80 nm thin sections, which complicates the interpretation of spatial relationships in three-dimensions (3D)^33^. Furthermore, TEM resolution is insufficient to fully capture the MC, approximately 300 nm in diameter and 500 nm in height, plus associated ciliogenesis membrane structures. In contrast, volume electron microscopy (vEM) enables the reconstruction of 3D models from a series of 2D images^34^. The application of vEM in life sciences has significantly enhanced our understanding of the intricate details and spatial relationships of structures at the cellular level^35^. Among vEM techniques, focused ion beam scanning electron microscopy (FIB-SEM) enables precise sectioning at intervals of ∼10 nm, providing isotropic resolution that minimizes distortion in 3D images^36^. This level of detail is particularly advantageous for visualizing cellular vesicles which can be as small as 30 nm in diameter^37^, as well as other structures generally below the resolution limits of most super-resolution light microscopy (SRM) techniques. Notably, FIB-SEM has been employed to describe membrane structures at the CV and sheath stages in ciliogenesis and in cilia resorption^16,38^, However, the characterization of MC docked pre-CV intermediates important in initiating ciliogenesis has not been defined by this approach.

In this study, we investigated the assembly of membranes on the MC during the early stages of ciliogenesis using FIB-SEM and SRM. We quantitatively investigated membrane docking and organization at the MC DAs and identified previously uncharacterized intermediate processes of ciliogenesis, including DAV docking and expansion, the organization of asymmetric tubular membranes, and a toroidal membrane structure preceding CV formation. By combining our vEM approach with genetic ablation studies we define ciliogenic roles for membrane trafficking regulators EHD1 and RAB8, and the DA protein CEP164, and the IFT-B complex protein IFT88. We found that EHD1, in association with MC-associated tubular membranes, biochemically interacts with CP110 and CEP97 and directs their removal from the MC to enable ciliogenesis to progress. Additionally, we find pre-CV tubular membrane assembly processes are linked to the recruitment of TZ proteins. This study demonstrates the application of isotropic vEM with quantitative structural analysis for understanding the molecular mechanisms of ciliogenesis.

## Results

### Identification of MC docked C-shaped membranes associated with ciliogenesis

To investigate the 3D organization of the MC and associated membranes during ciliogenesis by FIB-SEM, we employed a correlative light and electron microscopy (CLEM) approach with cells expressing the centriole/basal body (BB) marker GFP-Centrin 1 (GFP-CETN1) to precisely target regions of the cell for imaging. In RPE1 cells serum-starved for 3-6h to induce ciliogenesis cilia were detected in 32% of cells, while no cilia were observed in cells grown with serum (non-ciliating conditions) (Fig. 1a, b, Supplementary Table 1 and Supplementary Movie 1). A similar ciliation rate was noted in serum-starved primary human fibroblasts, with cilia present in 30% of cells imaged (Fig. 1b). Only a single CV covering the distal end of the MC was detected in both RPE1 and human fibroblast cells analyzed (Fig. 1b, c, Supplementary Fig. 1a, Supplementary Table 1 and Supplementary Movie 2). Strikingly, C-shaped tubulovesicular structures, defined as covering >50% of the circumference of the DA region but not fully encircling the MC (Fig. 1d, e), were present in approximately one-third of both cell lines, as well as in one serum-fed RPE1 cell (Fig. 1b, Supplementary Fig. 1b and Supplementary Movie 3). These findings suggest that the C-shaped membrane is a previously unrecognized intermediate structure in ciliogenesis that forms upstream of the CV stage.

**Fig. 1:**
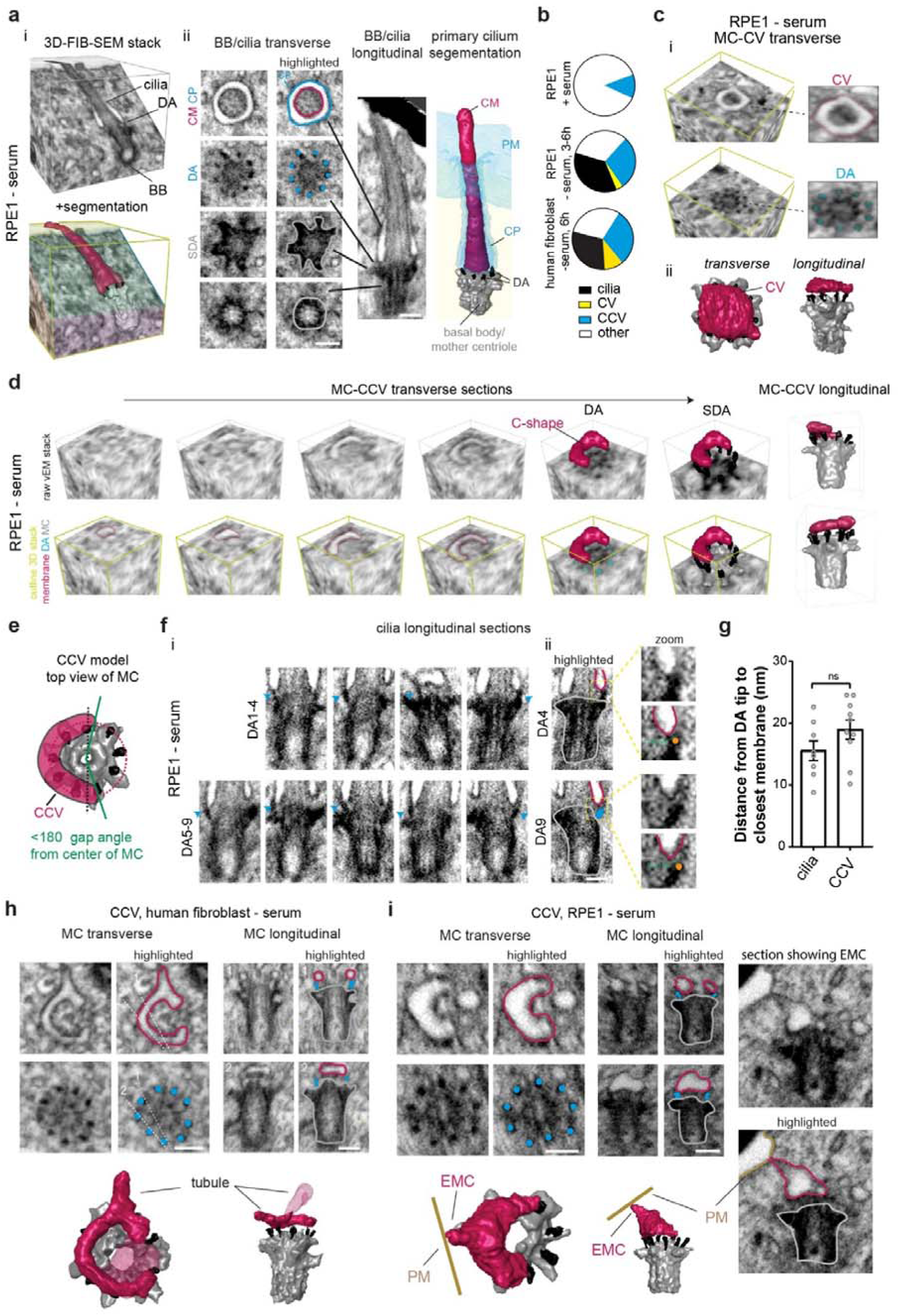
Identification of a CCV structure during ciliogenesis by vEM. **a** Representative FIB-SEM image stack and segmentation of primary cilium structure from a RPE1 GFP-CETN1 serum-starved (3 or 6h). i) 3D FIB-SEM image stack Segmented BB, DA and cilium (bottom image) from the volume integrated into the raw FIB-SEM image stack (top image). Different planar sections are colored differently in the bottom image. ii) Longitudinal and transverse sections showing CM: ciliary membrane (magenta), PM: plasma membrane, CP: ciliary pocket (outlined in cyan in vEM image). DAs are highlighted with cyan colored dots and MC/basal body is traced in grey in vEM images. PM and CP are shown in cyan in segmented image. Scale bars (white line): 200 nm. **b** Quantification of ciliary structures identified by FIB-SEM in serum-fed RPE1 cells (8 cells), serum-starved RPE1 (28 cells, 3 or 6h starved) and human fibroblast cells (10 cells, 6h starved). Other = not a CV, CCV or cilia. **c** i) MC transverse FIB-SEM 3D image stack containing a CV above the MC DA from a during human fibroblast cell described in **b**. ii) Longitudinal and transverse 3D segmentation of MC and CV from (i). **d** MC transverse FIB-SEM 3D image stack from a serum-starved RPE1 cell described in **b** with a C-shaped membrane above the DAs of the MC. 3D vEM planes (top and bottom images are identical positions) above the MC and at the MC DAs and SDA are shown. Top images showing mostly unobstructed membrane and MC structures present, bottom images highlight structures. Right images show 3D segmented structures for two different longitudinal views of the MC and C-shape membrane. **e** Model showing C-shaped MC docked membranes covering 50% or greater of the DA circumference are classified as CCVs. **f** Longitudinal sections showing DA-ciliary membrane interactions from cell described in **a**. (i) Images for all 9 individual DAs identified by a blue arrowhead. (ii) Highlighted DAs (cyan), BB (grey) ciliary membrane (magenta), and DA tip (orange dot). Magnified images (zoom) show representative 3D measurements for the closest position between the membrane and the end of the DA. **g** Plot showing distance from DA ends to membrane in cilia and CCV structures in RPE1 and human fibroblast cells. Cells were identified from 3 or more independent experiments (8 cells with cilia; 10 cells with CCV). Means ± SEM. The two-tailed t-test was not significantly (ns) different. **h, i** Representative 3D structural analyses of the CCV at the distal end of the MC in RPE1 (**i**) and human fibroblast (**h**) cells. Individual DA-membrane associations are shown as described in **f**. White dotted lines in **h** correspond to longitudinal sections of the centriole shown. Membrane tubules extending from the CCV **h** and an EMC connections to the PM **i** are shown. Scale bars (white line): 200 nm. Yellow lines are used in the vEM image stacks to highlight the 3D position of the section of the MC-associated structures shown (**a**, **c**, **d**).

To evaluate if the C-shaped membrane is associated with ciliogenesis on the MC, we determined the minimal distances between these membranes and the ends of the nearest DAs and compared these 3D spatial determinations to similar measurements performed on cilia (Fig. 1f and Supplementary Movie 4). The average minimal distance from the C-shaped membrane to the closest DA was found to be 19 ± 2 nm, which was not significantly different from that observed in cilia (Fig. 1g). These results support the conclusion that the C-shaped membrane is docked to the MC and is a ciliogenesis intermediate, which we refer to as the “C”-ciliary vesicle (CCV). Importantly, the CCV can be misidentified as DAV or CV stages of ciliogenesis depending on the specific longitudinal EM section of the MC being examined (Fig. 1h, i and Supplementary Fig.1b). The structure of the CCV suggests this membrane possesses tubulogenic membrane properties. This is further supported by observations that ∼50% of CCVs extend membrane tubules away from the MC (Fig. 1h, i and Supplementary Fig. 1b), with a third of these cells displaying tubular EMCs connecting the CCV to the PM (Fig. 1i and Supplementary Fig. 1b). The identification of CCV-EMCs indicates that membrane connections to the PM occur prior to the CV stage of ciliogenesis. Together, these findings reveal a previously unrecognized role for tubular membranes in cilium assembly and underscores the importance of employing vEM to elucidate the organization of MC-associated membranes during ciliogenesis.

### Characterization of DAV docking and organization in ciliogenesis progression

The discovery of CCVs and their potential misidentification as DAVs in 2D ultrastructure images raises questions about the organization and assembly of membranes on the DAs at earlier stages in ciliogenesis. To investigate if DAVs are associated with CCV assembly, we examined whether smaller membrane structures are present on the MC during ciliogenesis. The lower size limit of cellular vesicles is 30 nm in diameter, which can be resolved by FIB-SEM. Vesicles >30 nm were identified in FIB-SEM datasets after measuring their diameters and observing a continuous structure across two consecutive vEM sections. Membranes were considered docked to the MC if they were within 30 nm of the DA-ends (Fig. 2a, Supplementary Fig. 2 and Supplementary Movie 5), which corresponds to the average CCV-DA docking distance determined (Fig. 1g) plus an additional 10 nm to account for the resolution limit of FIB-SEM. MC docked vesicular/tubulovesicular membranes were defined as DAVs if a single structure covered less than 50% of circumference of the DA region (Fig. 2b). Our findings revealed that a quarter of RPE1 cells grown under serum-fed non-ciliating conditions did not have DAVs, while the remaining cells displayed DAVs or CCVs (Fig. 2c, d and Supplementary Table 1). In contrast, all serum-starved RPE1 cells had MC docked DAVs, CCVs or CVs (Fig. 2c, d, Supplementary Table 1). Similar results were observed in serum-starved human fibroblasts (Fig. 2d and Supplementary Table 1). Together these results confirm that DAV docking at the MC is associated with ciliogenesis progression.

**Fig. 2:**
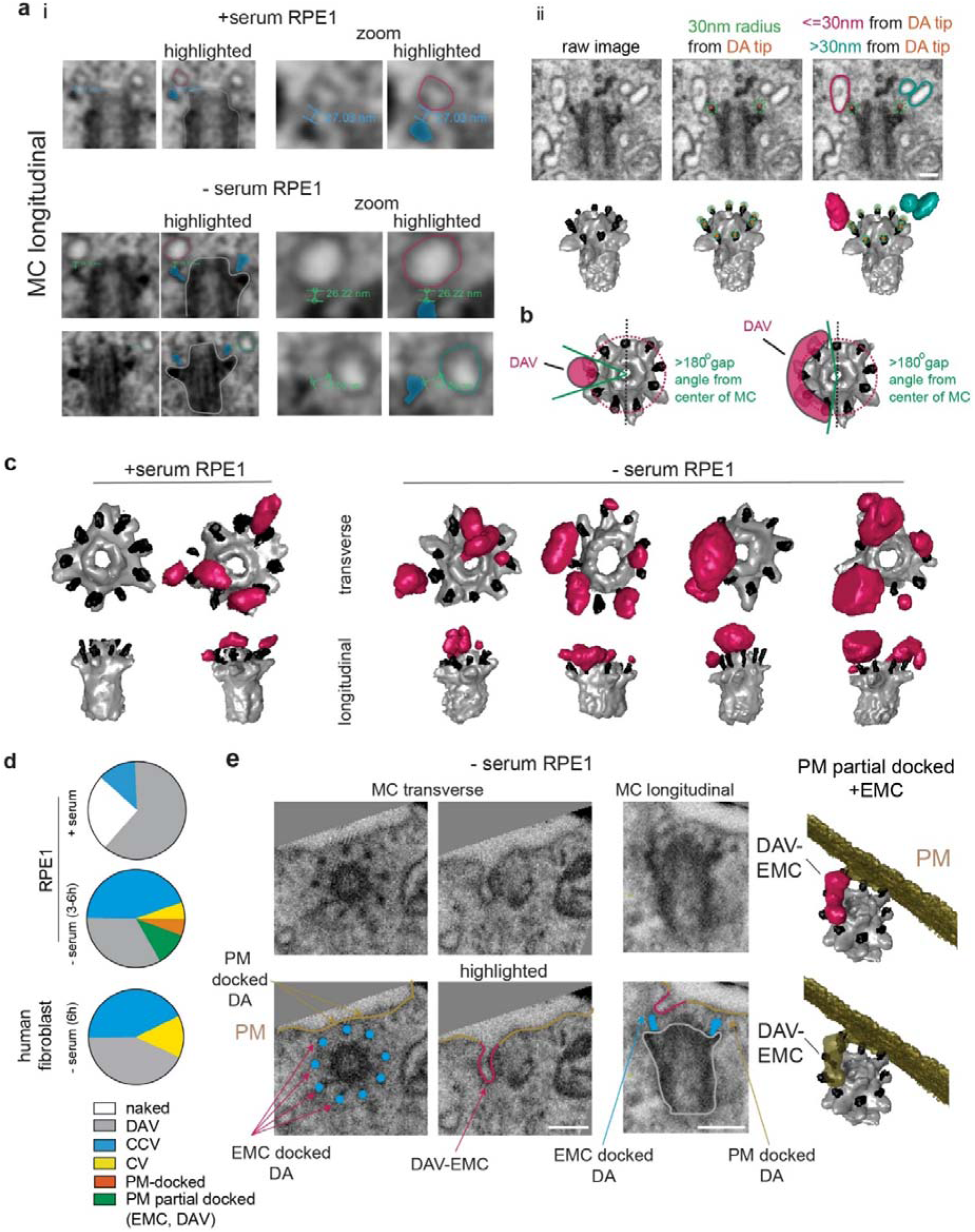
Characterization of DAV docking under ciliating and non-ciliating conditions. **a** i) FIB-SEM images showing membrane vesicle distances to the DA ends. ii) FIB-SEM and segmented images show a 30 nm spherical radius (green dotted line or mesh sphere) at the ends of the DA (orange dot). DAVs within 30 nm of the DA ends (magenta) and undocked vesicles more than 30 nm from the DA ends (cyan) are shown. **b** Model showing DAV covering less than 50% (>180° gap from center of MC) of circumference of the DA region. **c** Representative FIB-SEM segmentation of MCs from serum-fed (7 cells) or serum-starved (3-6h) (4 cells) RPE1 cells showing the absence (naked) and presence of DAV docked within 30 nm of the end of the DA. Segmentations for all DAV containing cells are shown in Supplementary Table 1. **d** Quantification of membrane structures observed in pre-axonemal MC in RPE1 (serum-fed, 8 cells; serum-starved, 18 cells) and human fibroblasts (- serum, 7 cells) shown in Supplementary Table 1. Cells imaged from 2 or more experiments. PM-docked corresponds to MCs where all DAs are docked to the PM without an axoneme protrusion into the membrane and a PM-partial docked example is shown in **e**. **e** vEM identification of partial MC docking to the PM docking via an EMC not associated with a CCV or later ciliogenesis stage. RPE1 cells were serum starved as described in **c**. EM images (*left panels*) and segmented images (*right panels*) show short EMC structures associated with a DAV stage ; referred to as DAV-EMC. Additional examples of MC with DAV-EMC MC are shown in Supplementary Table 1. Scale bars: 200 nm.

Interestingly, in two cells, the MCs were partially docked to the PM through EMC-like structures, while the other DAs were more than 30 nm away from the PM (Fig. 2d, e, and Supplementary Movie 6). In these cases, 2D EM images could lead to an incorrect interpretation that either the MC is not associated with the PM/ciliary membranes or that it is fully docked to the PM (Fig. 2e). Overall, our results indicate that both non-ciliating and ciliating cells typically contain DAVs within 30 nm of the MC DA ends, which we hypothesize are precursor membranes for the CCVs and are also capable of fusion with the PM.

To investigate the MC-DAV docking mechanism, we examined the effects of ablation of the DA protein CEP164, which has been reported to prevent membrane docking to the MC in TEM studies^39,40^. CEP164 was knocked out of RPE1 cells using CRISPR-Cas9 (Supplementary Fig. 3a,b) and we confirmed this proteins absence prevented cilia development, which was restored by expressing GFP-CEP164 (Supplementary Fig. 3a, c, d). Remarkably, FIB-SEM analysis of CEP164 KO cells identified DAVs docked to the MC in all cells examined (Fig. 3a and Supplementary Table 1). These findings suggest that CEP164 is dispensable for MC docking to ciliary associated membranes. Furthermore, a partially PM-docked MC with two DAVs was observed in one of the cells imaged (Fig. 3a and Supplementary Fig. 3e), which supports a model wherein fusion with PM can occur at the DAVs stage.

**Fig. 3:**
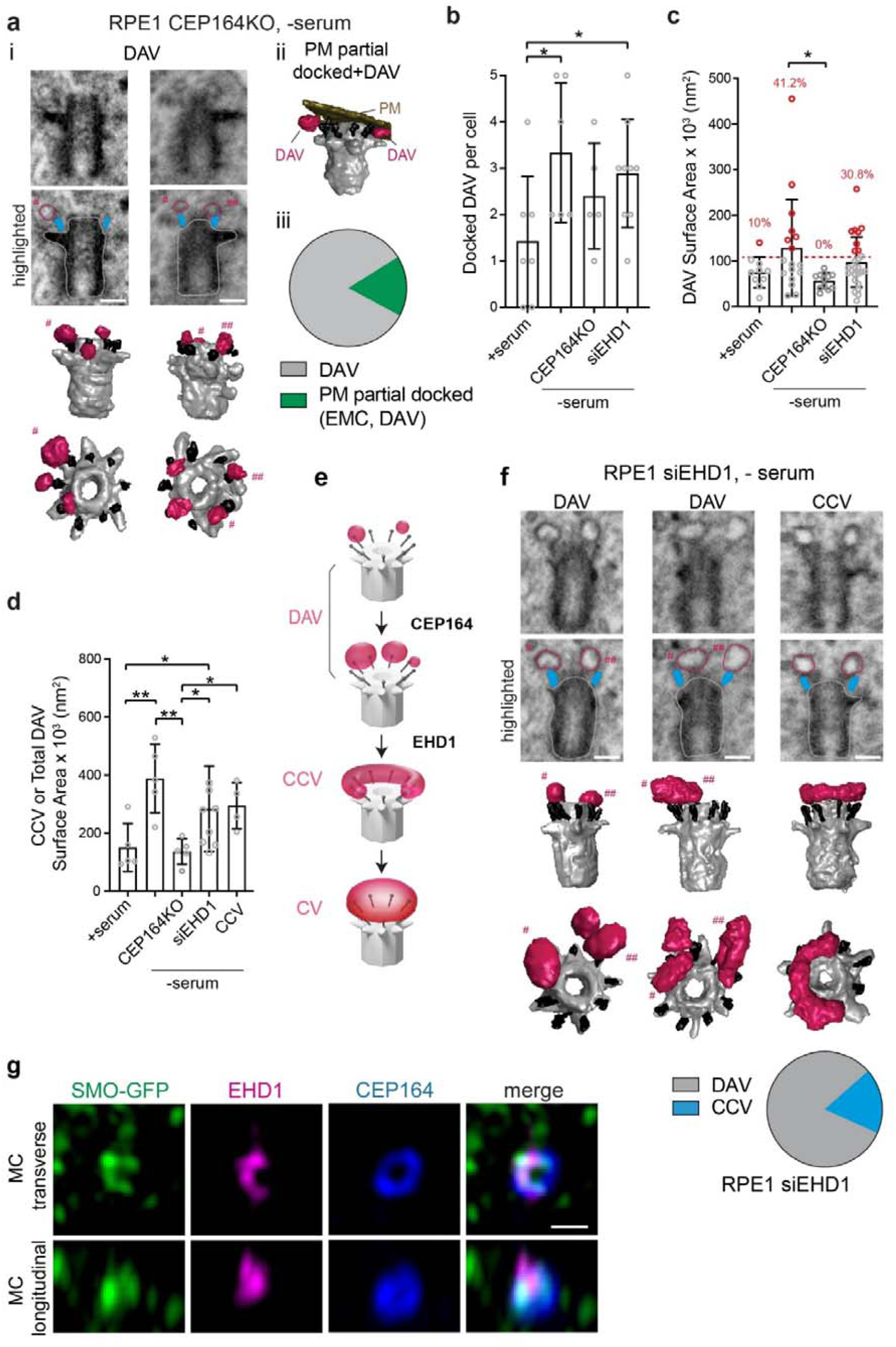
CEP164 and EHD1 function in ciliogenesis after DAV docking to the MC. **a** CEP164 knockout blocks ciliogenesis at the DAV stage. Representative segmented MCs (*top panel*) from CEP64 CRISPR KO cells (6 cells, shown in Supplementary Table 1) serum starved for 24h. (i) shows DAV stage, (ii) shows a MC partially docked to the PM with DAVs. Plot (*bottom panel*) shows quantification of MC structures detected in these cells. # and ## denotes specific DAVs shown. **b** Quantification of the total number of DAVs docked to individual MCs in RPE1 cells treated with serum (7 cells, described in (Fig. 2d)), and serum starved 3-6h (6 cells, described in (Fig. 2d)) or 24h (5 cells, CEP164 KO described in **a**; 9 cells,siEHD1 treated cells described in **e**. Mean ± SD, One-way ANOVA. * P<0.05. **c** Plot showing the individual DAVs docked to MCs in RPE1 cells treated with serum (10 DAVs, described in (Fig. 2d), serum starved 3-6h (17 DAVs, described in (Fig. 2d)), and serum starved 24h (12 DAVs CEP164 KO (**a**), 26 DAVs siEHD1 treated (**e**). Data points in red show DAVs with surface area 1 SD greater than the average surface area for serum treated cells (denoted by red dashed line). Mean ± SD. One-way ANOVA. * P<0.05. **d** Plot showing the total surface area of DAVs docked to each MC from RPE1 cells treated with serum (5 cells, described in (Fig. 2d), serum starved 3-6h (5 cells, described in (Fig. 2d), and serum starved 24h (5 cells CEP164 KO (**a**), 9 cells siEHD1 treated (**e**). Mean ± SD, One-way ANOVA. * P<0.05, *** P<0.001. **e** Model showing requirements for CEP164 and EHD1 in DAV expansion and CCV formation on the MC. **f** EHD1 depletion blocks ciliogenesis upstream of the CCV stage. Representative segmented MCs (*top panel*) from EHD1 siRNA treated (72h) cells (11 cells, shown in Supplementary Table 1) serum starved for the last 24h. Plot (*bottom*) shows quantification of MC structures detected in these cells. # and ## denotes specific DAVs shown. **g** SRM SIM image showing endogenous EHD1 localizes to asymmetric membrane structures along with SMO-GFP in RPE1 cells serum starved for 6h. DA are labeled with CEP164 antibody. MC transverse view shows a projection of image slices along Z-axis and MC longitudinal view shows a projection of image slices along X-axis. Scale bar: 500 nm.

To examine the ciliogenic potential of DAVs, we examined the distribution and organization of these membranes in RPE1 cells grown under non-ciliating and ciliating conditions and in starved cells lacking CEP164. Our analysis indicates that serum starvation promotes increased DAV docking to the MC compared to serum-fed wild-type cells (Fig. 3b). Notably, the average surface area of individual DAVs observed in wild-type cells under ciliating conditions was ∼125 x 10^3^ nm², which was ∼2-fold larger than in serum-fed cells (∼75 x 10^3^ nm²) (Fig. 3c). These results suggest that ciliogenesis progression is associated with bigger DAVs, which could result either from the docking of larger vesicles to the MC or from DAV fusion with smaller vesicles transported to the MC. The latter DAV expansion model is supported by observations that smaller DAVs coexist on the same MC with DAVs that are 3 to 4 times larger (Fig. 2c and Supplementary Table 1). Examination of the size of the DAVs in serum-starved CEP164 KO cells showed these membranes were typically less than 60×10^3^ nm², significantly smaller in size than found in wild-type starved RPE1 cells (Fig. 3c). These results indicate that CEP164 is required to organize larger DAVs on the MC during ciliogenesis. Notably, there is sufficient membrane in the DAVs of serum-starved RPE1 cells to form a CCV based on comparison of the surface areas of these membranes (Fig. 3d), but not in starved CEP164 KO RPE1 cells. Together these findings support a ciliogenesis progression model wherein larger DAVs are organized on the MC, a process requiring CEP164, followed by their fusion to form a CCV membrane structure (Fig. 3e).

### EHD1 functions in fusion of enlarged DAVs into the CCV

Because EHD1 has been linked to DAV functioning at pre-CV stages by TEM-based analysis^5^ and functions in membrane tubule formation and vesicle fusion^41,42,43^, we further investigated its ciliogenesis function by FIB-SEM in RPE1 cells treated with siRNA that depletes >90% of EHD1 protein^5^. All serum-starved EHD1-depleted cells displayed either DAVs or CCVs (Fig. 3f and Supplementary Table 1), with ∼80% displaying DAVs consistent with a role upstream of CCV assembly. EHD1 functioning at the CCV stage is further supported by structured illumination microscopy (SIM) observations of EHD1 on C-shaped structures that colocalized with SMO-GFP (Fig. 3g), a transmembrane receptor for the Hedgehog pathway which marks early ciliogenesis membranes on the MC^5,16^. EHD1-depleted serum-starved cells exhibited higher numbers of DAVs and had a greater total DAV surface area compared to cells grown under non-ciliating conditions (Fig. 3b, d), while the number and average size of these MC membranes in the EHD1 knockdown cells were comparable to serum-starved wild-type RPE1 cells (Fig. 3b, c). EHD1 function in CCV assembly on enlarged DAVs is further supported by the observation that the total amount of DAV membranes in EHD1-depleted cells were significantly higher than in CEP164 KO cells (Fig. 3d). Moreover, sufficient DAV membranes are present on the MC in EHD1-depleted cells to account for the size of an average CCV (Fig. 3d). Collectively, these results support a model in which EHD1 facilitates the organization of enlarged DAVs into the CCV during ciliogenesis (Fig. 3e).

### Assembly of the CV from DAVs and the CCV via a toroidal intermediate

The identification of the CCV as a prominent intermediate structure in ciliogenesis prompted further investigation into how these membranes are organized into a CV. Notably, FIB-SEM analysis revealed that ∼40% of cells with CCVs had associated DAVs in the gap between the ends of the CCV, that we refer to as the “C” gap (Fig. 4a, Supplementary Movie 7 and Supplementary Table 1). This finding raised the possibility that these DAVs fuse with the CCV to create larger structures with a reduced “C” gap (Fig. 4b). Indeed, a CCV expansion process is supported by comparison of the “C” gap in different cells (Fig. 4c). Notably, the lone CCV observed in serum-fed RPE1 cells showed the largest “C” gap at 153°, while the membrane gap in serum-starved cells was smaller ranging from 129° to 47° (average 93 ± 24°). Thus progressive fusion of the CCV with DAVs in the “C” gap could form to a continuous membrane around the MC upstream of CV organization.

**Fig. 4:**
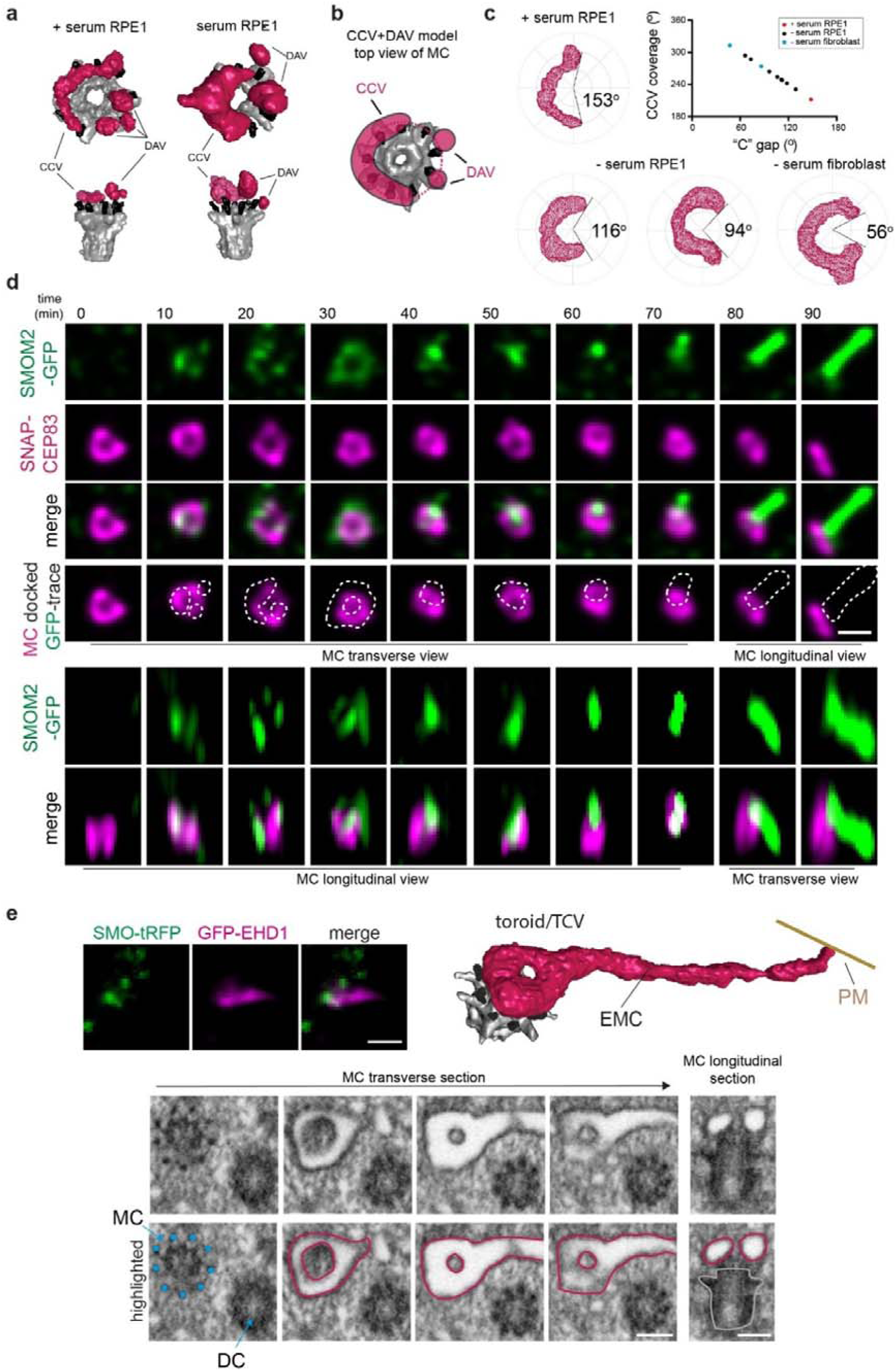
Ciliogenesis progression and a membrane toroid TCV stage. **a** Representative image of DAV docking in the CCV “C”-shaped gap. FIB-SEM segmentation from RPE1 cells shown in Supplementary Table 1. **b** Model showing DAV in CCV “C”-shaped gap on MC. **c** “C”-shaped gap angle determination from the center of the MC to the two ends of tubular CCV from RPE1 (9 cells) and human fibroblast (2 cells) cells as determined in Fig. 1e for cells shown in Fig. 2b. Plot shows distribution of CCV coverage and “C”-shaped gaps for each cell. ° = degree “C”-gap measured. **d** SRM SIM^2^ live cell time-lapse imaging of RPE1 cell showing ciliogenesis progression. Cells stably expressed SMOM2-GFP to monitor vesicle docking and ciliary membrane assembly at the MC marked by SNAP-CEP83 (labelled with SNAP-Cell647-SiR). Cells were imaged every 10 min following serum starvation. Images shown show pre- and post-SMOM2-GFP detection on MC DA. At the 70, 80, and 90 min time point the MC was sideways in the imaging plane. Panels with doted lines show the outline of SMOM2-GFP structures associated with the DA. MC transverse and longitudinal images correspond to planar position of the MC. Top and bottom images panels show projections of image slices associated with the MC in Z-axis and X-axis, respectively. Scale bar: 500 nm. **e** CLEM identification of a TCV structure in RPE1 cells expressing GFP-EHD1 and SMO-tRFP and serum starved for 6h. Cells were imaged by SD confocal (*top left panels*). FIB-SEM images (bottom panels) showing transverse and longitudinal sections of MC through the toroid membrane with and without traced DA (cyan), TCV associated membrane (magenta) and MC (grey). Additional FIB-SEM images are shown in Supplementary Fig. 5. Segmented image of the TCV with an EMC (top right panel). 6 total cells were imaged and analyzed; 1 TCV, 4 CV or short cilia, 1 naked MC. DC: daughter centriole. Scale bar for confocal: 2 µm. Scale bars for vEM: 200 nm.

To further investigate this ciliogenesis membrane assembly progression mechanism, we established an RPE1 cell line expressing the M2 variant of SMO (SMOM2-GFP), which labels pre-axonemal MC membranes^5^, and the DA protein CEP83 fused to a SNAP-tag. Using SRM-based SIM^2^ live-cell time-lapse imaging approaches, we could monitor membrane organization on the MC during ciliogenesis (Fig. 4d, Supplementary Fig.4 and Supplementary Movie 8). When the DAs faced upward or downward in the plane of imaging fluorescent-labelled SNAP-CEP83 appeared as a ring structure. Notably, SMOM2-GFP was detected accumulating as small punctate structures on one side of the DA-ring that progressed into CCV-like structures before consolidating into a more central elongating cilium structure at the end of the MC. Remarkably, prior to cilia growth the DA associated SMOM2-GFP shifted from an asymmetric DAV or CCV-like appearance to a ring-like structure within a 10 minute interval (Fig. 4d, Supplementary Fig. 4 and Supplementary Movie 8). These structures are consistent with either DAVs or CCV, but they may also suggest the presence of a toroidal membrane, resulting from DAV docking and fusion with the CCV in the “C” gap.

To investigate whether a toroidal intermediate is associated with ciliogenesis, we performed CLEM FIB-SEM on RPE1 cells expressing GFP-EHD1 and SMO-tRFP. These fluorescent reporters were used to screen for cells with potential tubular membrane intermediates at pre-axonemal growth stages on the MC. Strikingly, in one cell, we identified a membrane toroid docked to the DAs of the MC, which we designated the toroidal ciliary vesicle (TCV) (Fig. 4e, Supplementary Fig. 5 and Supplementary Movie 9). This TCV structure also displayed an EMC, consistent with PM connections being established at pre-CV membrane ciliogenesis stages. Together these observations support a ciliogenesis mechanism where the CCV organizes into a TCV upstream of the CV.

### Ciliogenesis tubular membrane organization directs MC uncapping and is associated with transition zone protein recruitment

We next investigated the association between ciliary membrane assembly intermediates and other ciliogenesis processes occurring at the MC, specifically MC uncapping and TZ recruitment. SIM SRM imaging in RPE1 and human fibroblasts cells expressing SMO-GFP suggests that CP110 removal advances with membrane accumulation on the MC (Supplementary Fig. 6). To further explore the relationship between ciliary membrane structure and MC uncapping, we conducted CLEM FIB-SEM imaging on ciliating RPE1 cells expressing GFP-CP110, SMO-tRFP and SNAP-CETN1. To enrich for early ciliogenesis membrane structures, cells were selected for FIB-SEM imaging that lacked elongated SMO-tRFP positive cilia and showed partial or complete GFP-CP110 loss from one of the SNAP-CETN1 positive centrioles (Fig. 5a). These CLEM studies revealed that the removal of GFP-CP110 from the MC is associated with progression from enlarged DAVs to the TCV stage, at which point CP110 is no longer present at the MC (Fig. 5a and Supplementary Fig. 7). Consistent with this model GFP-CP110 remained on the MC in serum-starved cells depleted of EHD1 that contained DAVs (Fig. 5b). To investigate the MC uncapping mechanism further we performed higher resolution SRM stimulated emission depletion (STED) imaging studies. Strikingly, we determined that GFP-CP110 removal from the MC primarily occurs in an asymmetric manner, with the remaining MC associated protein showing a “C” shape localization (Fig. 5c). A similar asymmetric uncapping process for CP110 and/or CEP97 was confirmed using SIM (Supplementary Fig. 8a, b) and expansion microscopy (ExM) (Fig. 5d, e). Together these findings suggest MC cap removal correlates with assembly of tubular membranes during ciliogenesis.

**Fig. 5:**
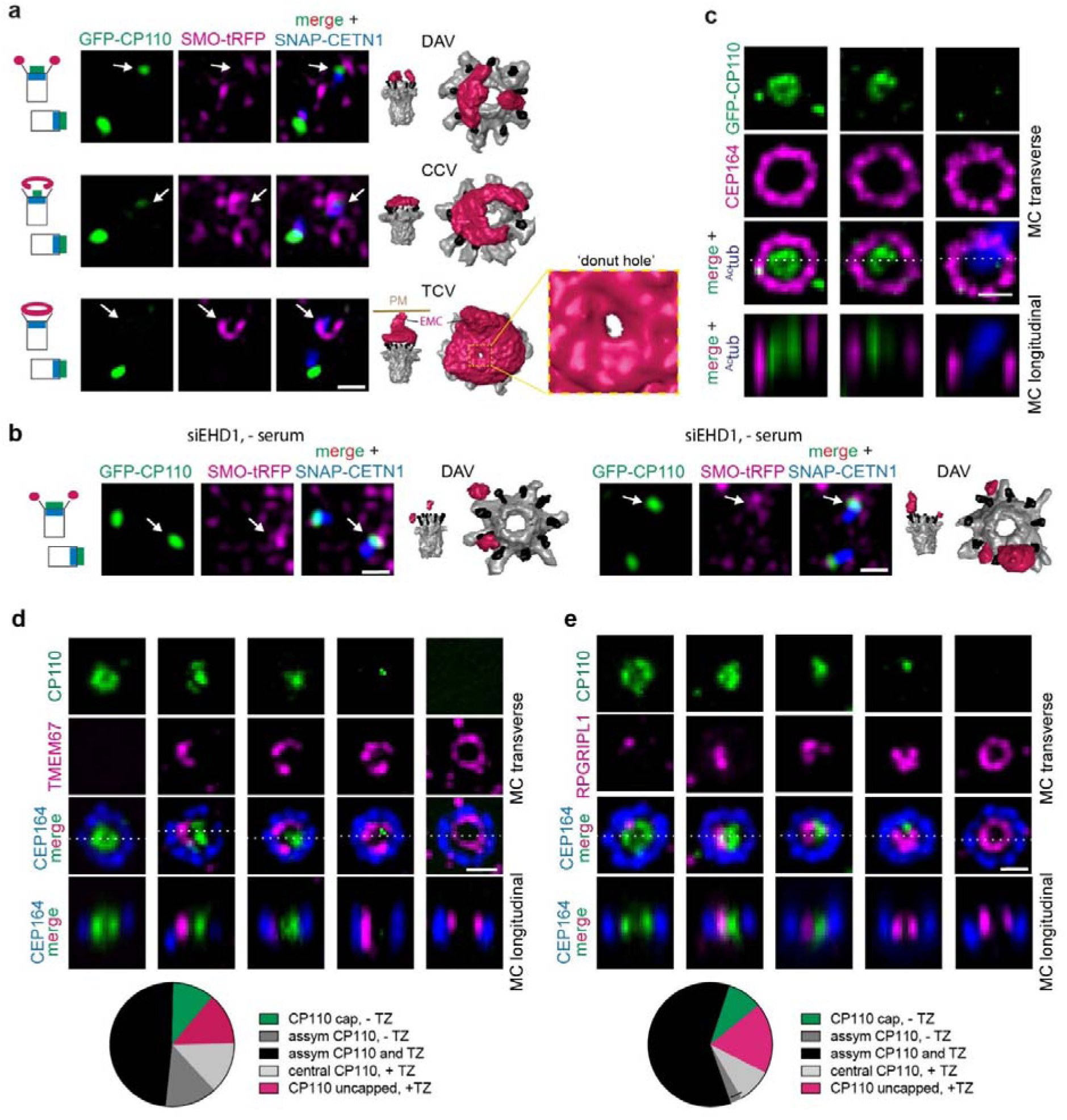
Asymmetric CP110 uncapping is associated with tubular membrane assembly and TZ protein recruitment on the MC. **a** DAV, CCV and toroid stages are associated with CP110 uncapping of the MC. RPE1 cells stably expressing GFP-CP110/SMO-tRFP/SNAP-CETN1 were serum starved for 6h. Cells were fixed and imaged by SIM (*left panels*) followed by CLEM FIB-SEM (6 cells) and segmentation (*right panels*). Zoomed region (*bottom panels*) shows ‘donut-hole’ region - see additional vEM images in Supplementary Fig. 7. Cartoons show fluorescence marker position on MC and DC associated with CP110 removal and DAV, CCV and TCV structures. White arrows show MC position. Additional segmented images are shown in Supplementary Table 1. Scale bar: 500 nm. **b** CLEM FIB-SEM analysis of CP110 capped MC and DC in EHD1 depleted cells. RPE1 cells described in **a** were treated with EHD1 siRNA for 72h with serum starvation for the last 24h. Cells were fixed and imaged by SIM (*left panels*) followed by CLEM FIB-SEM and segmentation (*right panels*). Cartoon as described in **a**. Scale bar: 1 μm. **c** SRM STED imaging demonstrates that CP110 removal from the MC occurs asymmetrically. RPE1 cells were serum starved for 6h and stained with CP110, CEP164, and ^Ac^tub antibodies and imaged by STED microscopy for the centriole markers and epifluorescence imaging for the cilia marker. Top images show a projection of images across the MC distal end region. Bottom panel show the orthogonal slice along the Z-axis corresponding the direction of the white line. Scale bar: 500 nm. **d** ExM showing asymmetric CP110 removal from the MC distal end of in relation to the TZ protein TMEM67. Top images show a projection of images across the MC distal end region. Bottom images show a slice along the orthogonal view along the Z-axis corresponding to the direction of the white line. Representative RPE1 cells serum starved for 6h. showing progression of MC uncapping (*left images*). Plot (right) shows the relationship between CP110 and TZ localizations observed for 37 cells from two independent experiments. Assym = asymmetrically distributed. Additional images showing reported localizations in the plot are included in Supplementary Fig. 8d. Scale bar 1 μm. **e** TZ protein RPGRIPL1 accumulates at the MC where CP110 has been removed. RPE1 cells were processed for ExM and stained with RPGRIPL1, CP110 and CEP164 antibodies and imaged and displayed as described in **d**. Plot showing CP110 and RPGRIPL1 MC distribution from two independent experiments for 33 cells as described determined in **d**. Additional mages showing reported localizations in the plot are included in Supplementary Fig. 8d. Scale bar = 1 μm.

Because MC uncapping is linked to TZ formation^44^ and TZ protein recruitment is associated with the CV stage^5,16^, we compared TZ and MC cap protein localization during ciliogenesis. Quantitative analysis of SRM ExM images revealed cells displaying a remarkable spatial correlation where the site of CP110 uncapping corresponds to the localization where TMEM67, a multispanning transmembrane TZ protein, and RPGRIPL1, a soluble TZ protein, accumulate on the MC (Fig. 5d, e and Supplementary Fig. 8d). Similar results were observed from SIM imaging studies of CP110 and CEP97 in RPE1 cells expressing the soluble TZ protein GFP-B9D2 (Supplementary Fig. 8a, b). Additionally, MCs with SMO-GFP donut-like structures observed by SIM imaging displayed TZ proteins CEP290 and TMEM67 accumulation either within or at the edge of the GFP-positive ‘donut-hole’, respectively (Supplementary Fig. 8c), consistent with the expected position of the TZ on the BB in ciliated cells. Based on these findings, we conclude that MC cap removal correlates with assembly of tubular CCV and TCV structures, and this process is coordinated with recruitment of TZ proteins associated with these membranes during ciliogenesis.

### EHD1 tubulogenic membrane properties direct MC uncapping

Based on the observation that MC uncapping correlates with ciliogenesis membrane structural organization, we hypothesized that proteins associated with these membranes may function in the removal of CP110 and CEP97 from the MC. To investigate this theory, we performed immunoprecipitation (IP) mass spectrometry (MS) studies using 293T cells expressing control LAP and LAP-CP110. Peptides for known CP110 interactors CEP97, the CP110 MC uncapping machinery HERC2 and NEURL4^31,45,46,47^, and CEP290^48^ were detected in the LAP-CP110 IP-MS analysis, but not in the LAP sample (Fig. 6a). Strikingly, peptides for EHD1 were specifically detected only in the LAP-CP110 samples (Fig. 6a). We further confirmed EHD1 interaction with CP110 in reciprocal IP-MS studies comparing cells expressing LAP-EHD1 and control LAP (Fig. 6b). Additionally, peptides for CEP97 and HERC2, a E3 ligase involved in ubiquitin-dependent degradation of CP110 during ciliogenesis^31^, were identified in LAP-EHD1 IP-MS samples. Immunoblot analysis of co-immunoprecipitation studies confirmed GFP-CP110 and GFP-CEP97 can interact with HA-tagged EHD1, but not with the other ciliogenesis membrane trafficking regulators tested (Fig. 6c, d). Comparison of EHD1 affinity for these MC cap proteins showed a stronger interaction with CP110 than CEP97 (Fig. 6e). Further biochemical characterization showed endogenous interaction between EHD1 and CP110 in 293T cells (Fig. 6f). To test if the EHD1-CP110 interaction is influenced by EHD1 association with tubular membrane structures, we conducted co-immunoprecipitation studies with mutant EHD1-K483E, which does not associate with tubular membranes^49^. Importantly, this mutant fails to rescue CP110 removal in EHD1-depleted cells localizes but localizes to the MC^5^. Strikingly, HA-tagged EHD1-K483E exhibited negligible binding to GFP-CP110 compared to the wild-type protein (Fig. 6g), indicating that EHD1 membrane tubulation function is essential for interaction with CP110. Based on these results we hypothesized that EHD1 at MC-associated tubular membranes controls CP110/CEP97 removal to promote ciliogenesis initiation. Moreover, this uncapping process may be directly regulated by increased trafficking of EHD1 to the MC observed in cells undergoing ciliogenesis^5^. To test this theory, we investigated the effects of GFP-EHD1 overexpression on ciliogenesis initiation in serum-fed cells. Strikingly, we found that nearly 70% of GFP-EHD1 transiently expressing RPE1 cells lost CP110 from the MC, compared to ∼30% of GFP alone expressing cells (Fig. 6h). Consistent with a role for EHD1 in pre-CV ciliogenesis initiating events only ∼20% of these GFP-EHD1 cells developed cilia (Fig. 6i). Interestingly, the number of cilia observed in GFP-EHD1 expressing cells was significantly higher than in cells expressing GFP (∼10%), suggesting EHD1 expression affects post-CV ciliogenesis progression. In contrast, the GFP-EHD1 K483E mutant did not significantly affect CP110 uncapping or ciliation compared to GFP expression (Fig. 6h, i). Collectively, these results indicate EHD1 on tubular CCV membranes directs MC uncapping during ciliogenesis via biochemical association with CP110 and CEPE97.

**Fig. 6:**
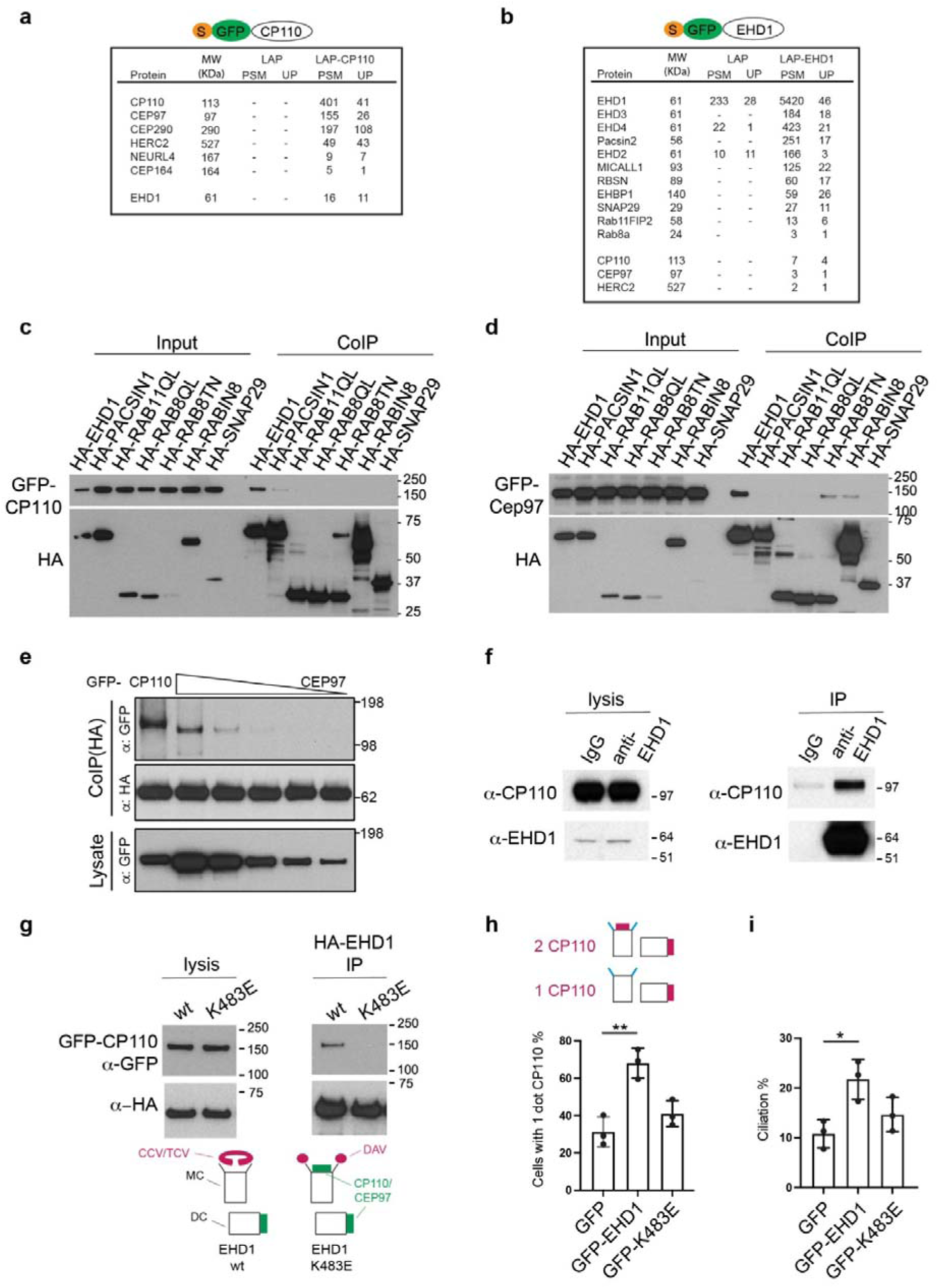
EHD1 promotes MC uncapping through its membrane tubulation function. **a** 293T cells stably expressing LAP (S-tag-TEV-GFP) and LAP-CP110 were immunoprecipitated and interacting proteins were identified by MS. **b** 293T cells stably expressing LAP, LAP-EHD1 were immunoprecipitated and interacting proteins were identified by MS. **c**, **d** EHD1 interacts specifically with CP110 and CEP97. Co-expression and co-immunoprecipitation of HA-tagged ciliogenesis membrane trafficking regulators with GFP-CP110 (**c**) and GFP-CEP97 (**d**) transiently transfected in 293T cells. GTP-locked RAB11A-Q70L and RAB8A-Q67L and GDP-locked RAB8A-T22N. **e** EHD1 binds more strongly with CP110 than CEP97. 293T cell lysate co-expressing HA-EHD1 and GFP-CP110 or GFP-CEP97 where immunoprecipitated with anti-HA antibodies. Cells were transfected with different amounts of GFP-CEP97 plasmid to examine MC capping binding properties. **f** Endogenous interaction between EHD1 and CP110. Immunoprecipitation was performed with EHD1 antibody or rabbit IgG as a control in 293T cells. Blots were probed with EHD1 and CP110 antibodies. **g** Immunoprecipitation studies demonstrate that wild type HA-EHD1 but not membrane tubule association deficient HA-EHD1-K483E mutants interact with GFP-CP110. Cartoon illustrates EHD1 wt and K483E associations with MC membrane docking and uncapping. Immunoprecipitation was performed with HA-beads. Proteins were detected using GFP and HA antibodies. **h**, **i** Wild-type EHD1, but not membrane tubulation deficient EHD1 K483E mutant promotes CP110 removal from the MC (**h**), and ciliation (**i**). RPE1 cells were transiently transfected with GFP, GFP-EHD1 or GFP-EHD1 K483E plasmids, and stained with the cilia marker ^Ac^tub and CP110 antibodies. Cartoon (**h**) shows conditions where CP110 is present (2 dots) or absent (1 dot) on the MC. Means ± SD (3 independent experiments, n: GFP=274 cells, GFP-EHD1=302 cells, GFP-K483E=282 cells), * P<0.05, **<0.01.

### RAB8 functions upstream of the membrane toroid structures in ciliogenesis

Genetic ablation studies demonstrated that RAB8 isoforms (RAB8A and RAB8B) are not important for MC uncapping in RPE1 cells, which is consistent with TEM results supporting a block in ciliogenesis at the CV stage^5^. However, DAV structures were also described^5^ raising the possibility that DAVs and/or CVs identified by TEM imaging could be misidentified tubular membrane intermediates. To clarify RAB8 ciliogenesis function, we performed FIB-SEM on serum-starved RPE1 cells described in Figure 5a after treatment with RAB8 siRNAs that inhibit ciliogenesis^5^. Cells lacking cilia were selected for FIB-SEM by CLEM using SMO-tRFP and SNAP-CETN1 markers, without screening for GFP-CP110 levels on the MC. Strikingly, our concerns about the use of TEM for precisely defining ciliogenesis mechanisms were further confirmed. Most RAB8 depleted cells showed arrested ciliogenesis at the CCV stage, while CVs and a single short cilium were detected in the other treated cells (Fig. 7a-e, Supplementary Fig. 9a, b and Supplementary Table 1). Remarkably, a CCV structure was discovered with a very small “C” gap (10°) that resembled an almost completed toroidal membrane (Fig. 7c), which supports the CCV to TCV assembly mechanism in ciliogenesis. These FIB-SEM analyses indicate that RAB8’s initial function in ciliary assembly is associated with CCVs, while stalled ciliogenesis at the CV stage supports a downstream requirement for RAB8 in axoneme extension. RAB8 functioning on tubular structures upstream of CV establishment is supported by the observation that RAB8 localizes with EHD1 on elongated membranes at the MC during ciliogenesis^16^. However, the RAB’s function in TCV formation appears to be unrelated to organizing membranes into tubular structures since RAB8 protein ablation does not affect CCV assembly. This conclusion is further supported by the fact that some RAB8 depleted cells showed an extended membrane tubule from the CCV and an EMC associated with a CV (Fig. 7c, e, Supplementary Fig.9a, b and Supplementary Table 1).

**Fig. 7:**
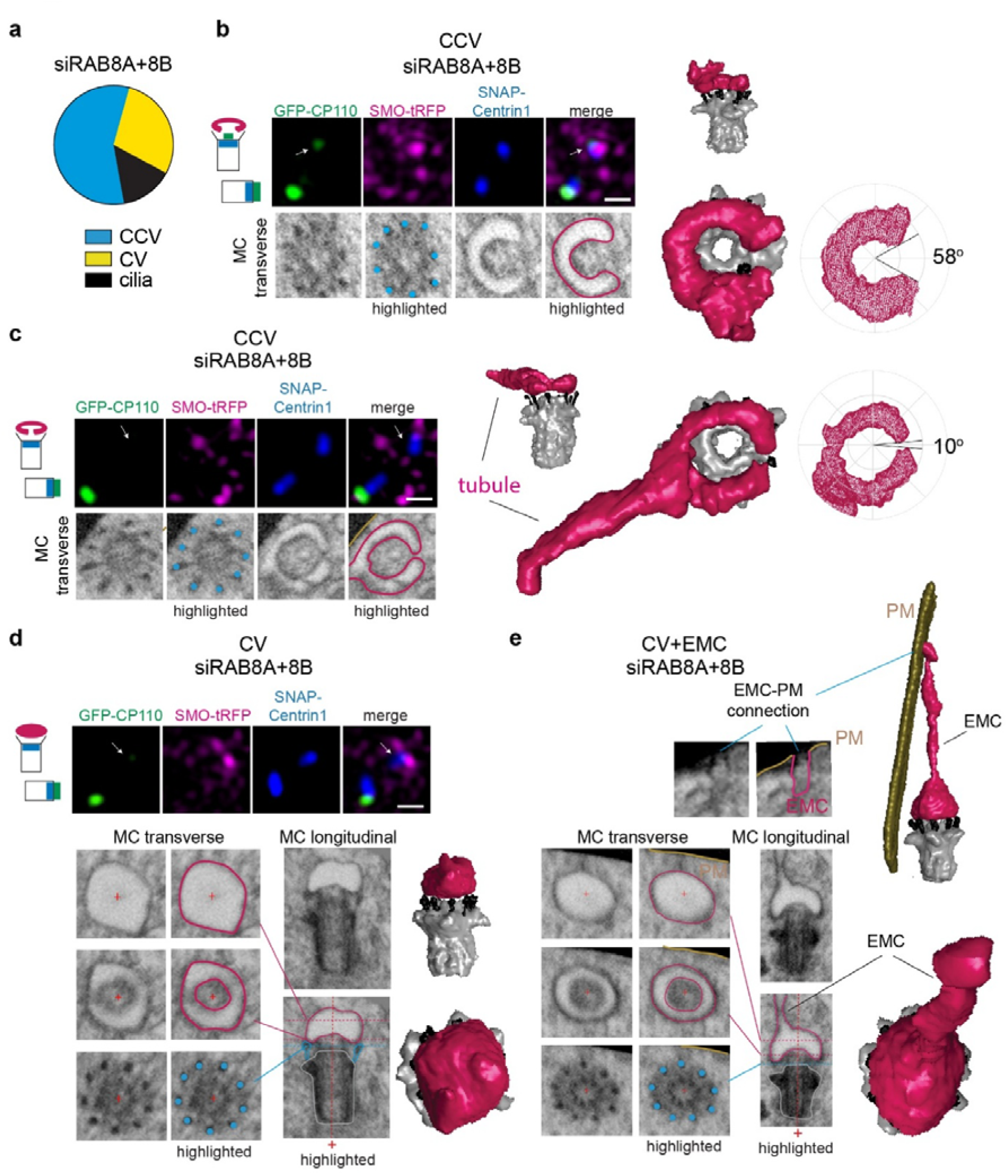
Rab8 functions in membrane toroid formation during ciliogenesis. **a** RAB8 requirements at the CCV to TCV stage in ciliogenesis. Plot shows ciliary structures identified in 7 RPE1 cells expressing GFP-CP110, SMO-tRFP and SNAP-CETN1 (shown in Supplementary Table 1) treated with siRNA that deplete RAB8A and RAB8B for 72h with serum starvation for the last 24h. **b**, **c** Immunofluorescence images of cells described in **a** showing GFP-CP110 removal from the MC (top panel immunofluorescence images). FIB-SEM images show transverse sections of MC with CCV structures (highlighted in magenta) on top of the MC DAs (highlighted with cyan dots). 3D segmented structures are shown along with the corresponding CCV gap determination map (right images). Cartoon show fluorescence marker position on MC and DC associated with CP110 removal and CCV structures **d** CV structure identified in RAB8 depleted cells described in **a** with complete CP110 uncapping from the MC. Longitudinal and transverse FIB-SEM images are shown with and without highlighted structures (DA cyan, CV magenta, MC grey) and the 3D CV structure reconstruction. Plus (+) is a positional marker for the transverse section of the MC shown. Cartoon as in **b** for CV structures. **e** Identification of a CV with EMC structure identified in RAB8A and RAB8B depleted cells described in **a**. vEM images and highlighted structures as described in **d** showing the EMC connection to the PM (EMC-PM). (+) as described in **d**. Scale bars for confocal images: 1 μm.

As expected, MC uncapping occurs in RAB8 depleted cells as GFP-CP110 was partially or completely removed from the MC in all RAB8 RNAi treated cells examined by CLEM SRM FIB-SEM (Fig. 7a-e). Notably, for the nearly completed toroidal membrane structure GFP-CP110 was undetectable (Fig. 7c), while in cells with larger “C” gaps some GFP-CP110 remained (Fig. 7b). These observations support requirements for tubular membrane assembly in instructing MC uncapping and indicate that RAB8-dependent CCV “C” gap closure upstream of TCV assembly is important for completing CP110 removal.

### IFT88 functions at pre-CV tubular membrane assembly stages during ciliogenesis

Roles for the IFT-B complex in ciliogenesis upstream of the CV stage are suggested based on the temporal accumulation of these proteins at the MC during ciliogenesis^5,13,16^. To investigate IFT-B complex pre-axonemal ciliogenesis function, we performed FIB-SEM imaging on IFT88 KO RPE1 cells which could not form cilia (Supplementary Fig. 10a-c), an effect rescued by the expression of mScarlet-IFT88. A critical IFT88 function in axoneme growth was confirmed by FIB-SEM analysis of 24h serum-starved IFT88 KO cells which revealed that ciliogenesis did not progress beyond MC docking to the PM (Fig. 8a, b). Strikingly, more than half of the cells imaged displayed ciliogenesis defects associated with earlier steps, particularly at the DAV stage (Fig. 8b, c) and one cell showed a TCV structure docked to the MC (Fig. 8b, d, and Supplementary Fig. 10d). These results suggest the kinetics of ciliary membrane development are affected prior to MC-docking to the PM linking IFT88 function to tubular membrane organization on the MC at pre-CV assembly stages of ciliogenesis.

**Fig. 8:**
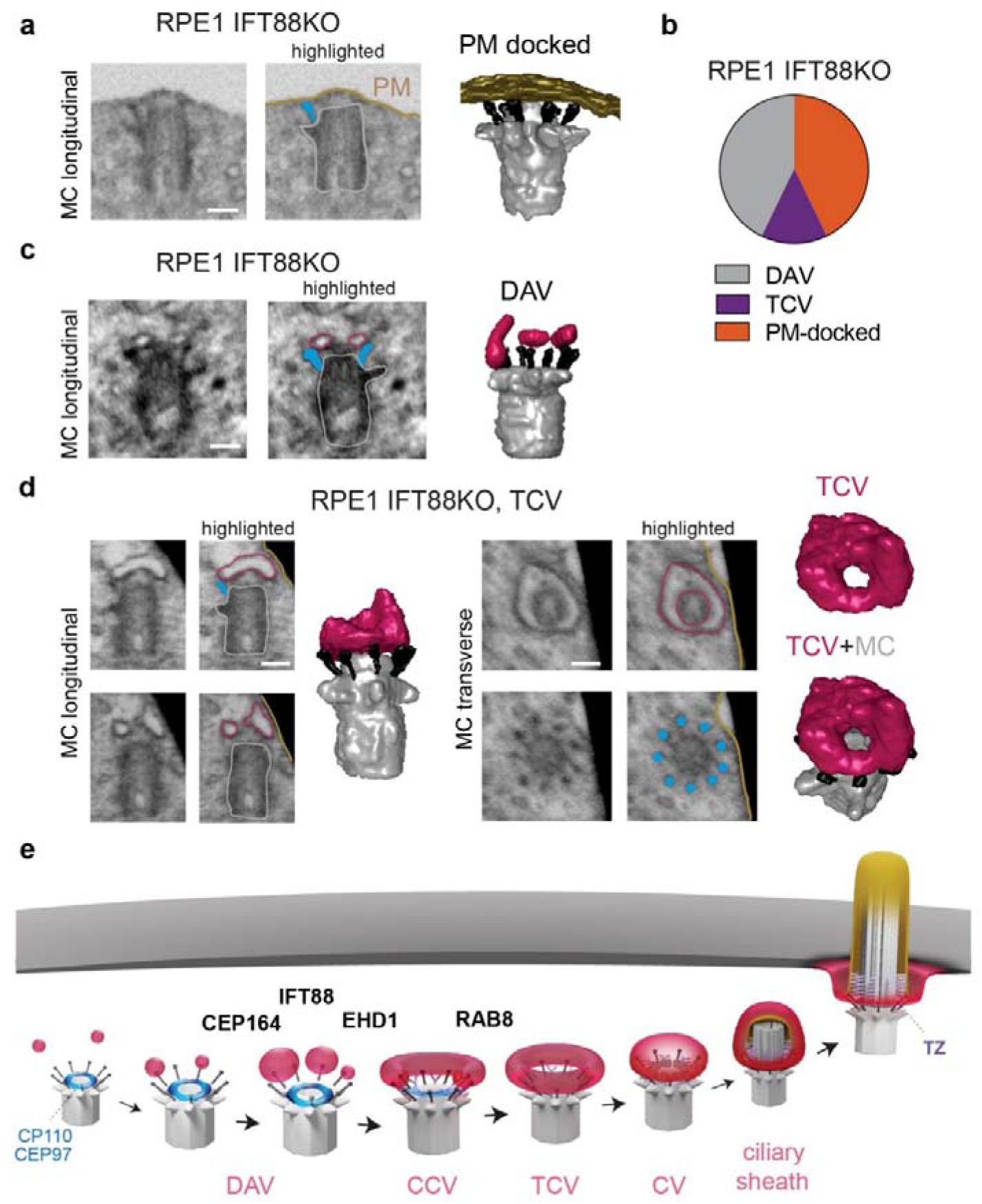
IFT88 is important at tubular membrane intermediate stages if ciliogenesis upstream of axoneme growth. **a**-**d** Characterization of ciliogenesis impairment in IFT88 KO cells expressing GFP-CENT1 by vEM. Ciliogenesis structures identified in RPE1 cells lacking IFT88 after 24h starvation. FIB-SEM and segmentation images showing representative PM-docked (**a**) and DAVs (**c**) and a TCV (**d**) from 7 cells with MC structures plotted in **b**. (**a**, **c**) FIB-SEM images show longitudinal. Highlighted structures shown on FIB-SEM images as described in Figure 1f. Scale bars 200 nm. **e** Schematic of membrane organization in intracellular ciliogenesis pathway.

## Discussion

Defects in membrane trafficking associated with ciliogenesis are causative in human ciliopathies^50,51,52,53^. The first description of membrane assembly at the MC distal end during ciliogenesis was reported over 60 years ago using TEM.^11^ However, the function and organization of these membranes during ciliary biogenesis has been enigmatic, limited by the investigative approaches available to researchers. The recent integration of vEM imaging approaches, specifically FIB-SEM, into the field of biology has unlocked the potential for directly visualizing cellular ultrastructure at low nanometer resolutions^35,36^. Only a few studies have employed quantitative analysis of cellular structures by vEM^54,55,56,57^, and here we show this approach can be used to understand organelle biogenesis dynamics in the low nanoscale range. Using this and other advanced imaging approaches we reveal the intricate steps of MC-associated vesicle docking, fusion and reshaping, along with establishing new mechanistic connections to MC uncapping and TZ protein recruitment function in ciliogenesis (Fig. 8e).

Using FIB-SEM, we show it is possible to spatially resolve ciliogenesis-associated structures isotropically at 10 nm resolution. This enabled the quantitative characterization of different ciliogenesis membrane intermediates through examining their frequencies, sizes, and distributions on the MC. Our findings support a mechanism in which increased membrane docking of DAVs to the MC occurs during ciliogenesis, followed by the expansion of DAVs, which we link to the DA protein CEP164. The membrane shaping and fusion regulator EHD1 was found to function in the formation of tubular CCV membranes from enlarged DAVs. Interestingly, vEM studies on differentiating mouse granule cells showed tubular membranes associated with the MC attributed to resorption of cilia, suggesting a similar membrane organization process could function in both assembly and disassembly of the cilia^38^. Our vEM analysis also redefined RAB8 and IFT88 ciliogenesis functioning in association with assembly of tubular membrane intermediates. Specifically, our results suggest that RAB8 is involved in CCV closure to a TCV, possibly through regulating membrane fusion. Our finding that IFT88 is also important for efficient progression of ciliogenesis upstream of ciliary axoneme growth merits additional investigation. This is consistent with the presence of the IFT-B complex at the centrosomes in non-ciliating cells^58,59^. Notably IFT-B components and have been linked to pre-axoneme ciliogenesis processes in multiciliated cells (MCCs)^60^, although it remains unclear if MCCs cells utilize a similar ciliary assembly mechanism as primary ciliated cells. Notably, FIB-SEM imaging in Rab34 knock-out RPE1 cells^25^ suggests a role for this membrane trafficking regulator upstream of CV assembly, although comparison of docking and vesicle size parameters will be necessary to more precisely resolve this proteins role in the ciliogenesis as specified in this study.

This study also provides new insights into the connection between ciliary membrane formation and MC to BB transition, specifically through the removal of the CP110/CEP97 MC cap. Our findings suggest that EHD1 can form a complex with CP110 and CEP97, facilitating the removal of these proteins from the MC through its membrane tubulation associated function. One potential mechanism for this coordination involves the recruitment of HERC2 to the MC by EHD1, which promotes the ubiquitination of CP110^31^. Interestingly, CP110 has been shown to inhibit the function of CEP290 in TZ formation^48^. This aligns with our observations that TZ proteins accumulate only where CP110 and CEP97 have been removed from the MC distal end. We propose that the TCV ciliogenesis intermediate in association with TZ proteins, serve as a ciliogenesis progression checkpoint prior to the initiation of axoneme growth and ciliary extension.

The mechanisms governing the assembly and functionalization of the multiprotein transition zone (TZ) complex, comprising soluble and transmembrane proteins essential for regulating ciliary transport, remains poorly understood. This knowledge gap is highly significant as TZ proteins are frequently mutated in human ciliopathy^1,2^. Our finding showing the transmembrane TZ protein TMEM67 on CCV-like structures suggests this protein is recruited on pre-ciliary membranes targeted to the MC. However, it is also possible EMCs mediate TMEM67 trafficking to MC associated membranes, as well as other TZ proteins and ciliogenesis factors. Importantly, our findings supports a model where EMCs mediate MC connections to the PM^16^, forming from CCV and possibly earlier DAV membranes. What is clear is that TZ protein accumulation at the MC coincides with CCV and TCV assemble suggestive that the TZ is being organized. Interestingly, the surface of CCV and TCV membranes and the cilium membrane both face the MC DAs and likely share a similar curvature, and in the case of the TCV encircles the MC. Thus it is conceivable that these tubular ciliogenesis membranes may be organized similarly to the mature ciliary structure, complete with a TZ capable of regulating the targeting of proteins critical for ciliary growth, including axonemal assembly factors such as molecular motors, microtubule-associated proteins (MAPs) and EB1/3^61,62^. It will be interesting to determine whether the donut-shaped membrane shares a similar phospholipid composition with the ciliary membrane, which in the cilia is regulated by cilia-enriched INPP5E that converts PtdIns(4,5)P2 into PtdIns(4)P^63^. Notably, within the cilium, the membrane exhibits negative curvature along its length, transitioning to positive curvature at its base into the cytoplasmic face of the ciliary pocket (CP) membrane. Ciliogenic EHD and PACSIN family proteins, known to be associated with positive-curvature membranes^64,65^, localize to the CP, but not within the cilia^16^, This further suggests a partitioning mechanism by the TZ at the CCV/TCV stages that helps facilitate the establishment of the negatively curved ciliary membrane as ciliogenesis progresses.

To our knowledge, naturally occurring toroidal membrane vesicles have not been documented in cells. Modeling predictions suggest that shaping membrane tubules into C-shaped and toroidal structures is highly energetically unfavorable^66^, particularly for smaller membranes such as those observed at the MC during ciliogenesis. Notably, the final fusion of the C-shape ends to form the toroid is the highest barrier energy state in these models. In ciliogenesis, organization of these tubular structures is likely aided by anchoring membranes to the DAs through protein-protein interactions. These MC-membrane connections likely reduce the energy barriers affecting the stability and formation of tubular structures, thereby reducing the energy burden associated with membrane bending. Additionally, structural support for membrane curvature may be enhanced by the establishment of TZ, which features Y-linker structures that extend from the membrane to the central region of the MC. Our findings suggest a compelling role for RAB8 in balancing bending energy and the release of osmotic potential energy during the closure step of CCV to form TCV. However, RAB8 function could simply be associated with regulating final DAV docking in the “C”-shape gap or in their fusion with the CCV to generate the TCV. RAB8 function on CCVs could also be associated with the TZ, as it is known to interact biochemically with the core TZ protein CEP290^48^. Overall, our work indicates roles for proteins on the membrane and at the distal end of the MC in establishing the membrane curvature of CCV and TCV structures.

Lastly, our findings shed new light on the MC-membrane docking regulation and ciliogenesis initiation. Previous ciliogenesis models proposed DA proteins function in the initial membrane docking to the MC^10,13,14^, recruitment of the kinase TTBK2 for MC uncapping^67,68^. and in RAB8-positive membrane growth of ciliary membrane in concert with the axoneme formation^9,39,69,70^. Interestingly, our detailed analysis of the spatial dynamics of vesicle docking and the size of these membranes reveals that small vesicles can dock to the DAs without facilitating ciliogenesis. Thus, progression to ciliation is in fact triggered by the fusion of DAVs to form tubular membranes which encircle the MC via processes requiring CEP164 and EHD1. The observed partial MC docking to the PM in cells lacking CEP164 suggests that this protein plays a previously unrecognized role in ciliary membrane organization, potentially through the recruitment of membrane fusion and shaping factors. These findings could have significant implications for understanding human ciliopathies associated with CEP164^51,71^.

In this study, we have achieved a more comprehensive understanding of the spatial and temporal control of ciliary membrane assembly at the MC (Fig. 8e). The advanced imaging approaches employed permitted precise identification of requirements for membrane structure organization and key regulators in ciliogenesis, including MC DA proteins and capping proteins and membrane trafficking regulators and the IFT-B complex. We expect this work will prompt new investigations aimed at further characterizing the function of these and other ciliogenesis factors, informed by the architectural model presented. This study highlights the immense potential of utilizing vEM techniques and quantitative 3D structural analysis to unravel intricate isotropic complexities associated with the biogenesis of cilia and other organelles.

## Methods

### Antibodies and reagents

Commercial antibodies used were as follows: anti-Acetylated tubulin (Actub, clone 6-11B-1, 1/10000, T6793, Sigma), β-Actin−Peroxidase antibody (clone AC-15, 1/30000, A3854, Sigma), anti-EHD1 (EPR4954, 1/500, ab109311, Novus Biologicals), anti-RPGRIP1L (1/200, 55160-1-AP, Proteintech), anti-TMEM67 (1/200, 13975-1-AP, Proteintech), Rabbit anti-CEP164 (1/500, 22227-1-AP, Proteintech), chicken anti-CEP164^16^, Rabbit anti-CP110 (1/1000, 12780-1-AP, Proteintech), mouse anti-CP110 (1/500, MABT1354, Millipore Sigma), anti-CEP97 (1/1000, A301-945A, Bethyl), anti-GFP Alexa 488 (1/1000, A21311, Molecular Probes Life Technologies), DAPI (1/2000, 62248, Thermo Scientific), Hoechst (1/3000, H3570, Molecular Probes Life Technologies) Rabbit anti-Pericentrin (1/2000, NB100-61071, Novus) and Mouse anti-Arl13b (1/500, N295B/66, NeuroMab) and all Alexa Fluor Dyes conjugated secondary antibodies were from Life Technologies. Goat anti-chicken IgY CF640R (Biotium, 20084), goat anti-rabbit CF568 (Biotium, 20099) and goat anti-mouse CF488 (Biotium, 20010) were used for ExM staining. SNAP-Cell647-SiR reagent was purchased from New England Biolabs. Atto594 and Atto647N conjugated secondary antibodies for STED imaging were from Millipore-Sigma. Doxycycline hydrochloride was obtained from Sigma and used according to manufacturer’s instruction.

## Cell lines

Human hTERT-RPE1 (CRL-4000) and 293T (CRL-3216) cell lines were obtained from ATCC. Human fibroblast cell line was obtained and cultured as previously described^72^. LAP, LAP-EHD1, GFP-CETN1, SMO-GFP, SMOM2-GFP, GFP-B9D2, SMO-tRFP and LAP-EHD1+SMO-tRFP RPE1 cell lines were generated using lentivirus or Flp-In system as previously described^5,16,73^. The SMOM2-GFP+SNAP-CEP83 cell line was generated by introducing SNAP-CEP83 lentivirus into the SMOM2-GFP cell line. For SIM FIB-SEM CLEM studies, RPE1 cells were infected with LAP-CP110, SMO-tRFP^5^, and SNAP-CETN1^16^ viruses, then isolated as single-cell clones. Human fibroblasts were infected with SMO-GFP lentivirus to make the stable cell line. 293T cells expressing LAP and LAP-EHD1 or LAP-CP110 were stably generated using lentiviral transduction^73^. DNA transfections were carried out using X-tremeGENE 9 (Roche). LAP, LAP-EHD1 and LAP-CP110 expression was induced with 0.5 μg/mL doxycycline^5^. For knockdown experiments, cells were transfected with siRNA duplexes (Dharmacon) using RNAiMAX (Invitrogen) according to manufacturer’s instruction and were fixed for analysis after 72□h treatment.

### Immunofluorescence

To promote ciliogenesis, cell lines were serum-starved for 3, 6 or 24 h. After serum starvation, cells were fixed using 4% paraformaldehyde or cold methanol for 10 min. Blocking was performed for 10 min with 1% BSA in PBS 0.1% Triton X-100 or Saponin, followed by immunostaining in blocking solution with indicated antibodies in figure legends. Confocal images were taken using a Marianas spinning disc confocal microscope (Intelligent Imaging Innovations) equipped with a 63× 1.4 NA as indicated in figure legends and a CMOS camera (ORCA-Fusion, Hamamatsu). Cellular labeling was done using SNAP-Cell647-SiR (New England Biolabs) substrate according to the manufacturer’s recommendation and as described in figure legends. Ciliogenesis and CP110 uncapping were quantified as described^5^.

### Structural illumination microscopy (SIM)

The coverglass with fixed cells was incubated with 100 nm Tetra Speck beads from Life Technologies in the final wash, were mounted onto glass slides using ProLong Glass Antifade Mountant from ThermoFisher. 3D-SIM imaging was conducted using a Nikon N-SIM microscope, a GE DeltaVision OMX SR imaging system, or a Zeiss Elyra 7 SIM microscope, as indicated in the Figure legend. Specifically, Nikon SIM images were captured with a SR Apo TIRF x 100/1.49 NA oil immersion objective from Nikon and an EMCCD camera (Andor DU-897E), while Zeiss SIM images were obtained with a 63 x 1.4 NA oil objective from Zeiss and a PCO edge sCMOS camera, as previously described. The image stacks were acquired at a z-distance of 0.1 μm, and alignment parameters for all color channels were carefully determined during the calibration process using Tetra Speck beads. The Nikon Elements or Zeiss Zen software was utilized for image reconstruction and processing, and tiffs were edited using FIJI. Intensity profiles were generated following established protocols.

### STED microscopy

STED imaging utilized the Leica TCS SP8 STED 3X system, which is equipped with a white light laser for excitation. The imaging employed a ×100 oil-immersion objective from Leica with a numerical aperture (N.A.) of 1.4. For dual color STED imaging, Atto594 and Atto647 congregated secondary antibodies were used for imaging.

### Expansion microscopy (ExM)

ExM samples were prepared as previously described with modifications^74^. Cells grown on coverslips were fixed and subjected to immunofluorescence staining with primary and secondary antibodies as described above. The coverslips were then incubated with AcX solution for 18 h and subjected to gelation (8.6% Sodium acrylate, 20% acrylamide, 0.15% N,N′ -Methylenebisacrylamide, 2M NaCl in 1x PBS supplemented with 0.2% APS and 0.2% v/v TEMED). The gel was digested with proteinase K in digestion buffer (0.5% Trition X-100, 25 mM EDTA, 800 mM NaCl, 50 mM Tris) for 18 h and stained with DAPI before being placed in ddH2O. After expansion, images were captured with Yokogawa W1 spinning disk on a Leica DMi8 microscope using a 63x NA1.2 water immersion lens. Images were acquired with the MetaMorph® Microscopy Automation and Image Analysis Software (Molecular Devices) and processed using Fiji.

### Time lapse SIM microscopy

Super-resolution SIM live-cell imaging was performed on a Zeiss Elyra 7 SIM^2^ system, with the environmental chamber maintaining a temperature of 37°C supplemented with 5% CO2. To conduct Apotome SIM time-lapse experiments, multiple positions were defined within the position window of the Zen 3 Black software. Images were captured using a 40x 1.4NA oil objective, with three phases obtained at each time point every 10 min for 18 h. To capture the MC/BB movement during ciliogenesis, a z-stack of approximately 3 µm was captured using Leap mode for each time point. Figures display projections of 3-5 slices from the stack around the MC/BB.

### CRISPR Cas9 knockout cell lines

peSpCas9(1.1)-2×sgRNA and peSpCas9(1.1)-2×sgRNA-(IFT88, donor) plasmids used for IFT88 and CEP164 knockout were gifts from Kazuhisa Nakayama (Addgene plasmid #80769 and #80768). CEP164 gRNA was designed as described^40^ and cloned into peSpCas9(1.1)-2×sgRNA. IFT88 and CEP164 knockout RPE1 cell lines were generated as described^75^ Rescue experiments were performed in stable cell lines expressing GFP-CEP164 or mScarlet-IFT88.

### CLEM sample preparation

Sample preparation was performed as described previously^16^. Briefly, cells were grown on alphanumerically coded gridded coverslips. After fixing, various cells of interest were imaged by spinning disc confocal microscopy as described above. Immediately after fluorescence imaging, bright field images of the gridded pattern containing the cells were acquired to generate an accurate “target map” of candidate cells for interrogation by FIB-SEM. The cell samples were then post-fixed, stained, dehydrated, and embedded in PolyBed resin according to standard protocols. This allowed the etched alphanumeric pattern to be transferred to the resin. The blocks were then gently cleaned, affixed to an SEM stub with conductive silver paint, and sputter coated with a thin conductive layer of carbon before transfer to the FIB-SEM instrument. For SIM/FIB-SEM, RPE1 cells expressing were grown in 35 mm glass-bottom dishes with alphanumerically coded gridded coverslips. The cells were serum-starved for 3 h or transfected with EHD1 or RAB8a+RAB8b siRNA for 48 h and serum-starved for an additional 24h before fixation. Then cells were fixed with 4% paraformaldehyde and 0.25% glutaraldehyde in 0.1 M sodium cacodylate buffer for 30 mins at RT and washed with 0.1 M sodium cacodylate buffer 3 times before imaging. 3D SIM images were collected on a Nikon N-SIM with a 100x objective. Phase contrast images of the target cells and the alphanumerical pattern of the coverslip were taken with both the 100x objective and a 10x objective. The alphanumerical pattern will be transferred to the resin block after embedding and the target cell can be relocated through the pattern under FIB-SEM.

### FIB-SEM imaging

FIB-SEM imaging was performed in a Zeiss Crossbeam 550 (Carl Zeiss Inc.) in conjunction with ATLAS3D software (Fibics Inc.), as previously published^16^ with a few modifications. Briefly, a platinum and carbon patterned protective pad was deposited with the FIB operated at 700□pA, and data collection was executed with the FIB and SEM operated simultaneously. The FIB was operated at 30□kV, 700□pA, SEM operated at 1.5□kV, 1□nA, and back scatter signal was recorded at the in-column EsB detector operated with a 900□V grid voltage. The “ROI” images were acquired at 3□nm pixel sampling and 9□nm milling increments, with total dwell time of 3□µs per pixel. An imaging run covering portions of a cell typically lasted ∼20□h and generated a stack of ∼1000 high resolution images; however, the volume containing the centriolar area was much smaller. These images were registered using in-house IMOD based scripts, and subsequently cropped, binned and inverted to yield registered, isotropic (9□×□9□×□9□nm) volumes in mrc format. These data provided a high-resolution vEM readout corresponding to the targeted cellular features imaged previously by fluorescence, and centrioles could be easily identified without further correlative fiducial markers.

### FIB-SEM segmentation

FIB-SEM reconstructions were analyzed using Dragonfly or 3D slicer software. 3D volume segmentation models were generated using Dragonfly with ciliary-associated structures (centrioles, ciliary membrane, PM, and tubules) by using a combination of automatic thresholding and manual assignments. To ensure accuracy, segmentation assignments were verified across all three (xyz) FIB-SEM image planes. Additionally, the segmentation of structures was independently performed by 3-5 individuals for validation.

### vEM FIB-SEM image analysis

CCVs gap determination was performed on Dragonfly segmentations that were exported as contour meshes and subsequently converted to NumPy arrays of cylindrical coordinates. This array was sorted by ascending θ with the largest difference in θ taken as the C-Shape Gap. CCV were characterized as having membrane gaps less than 180°. Membrane distances to DA distal end in cilia, CV, and CCV was measured in 3D for vEM stacks using the Dragonfly ruler tool. For CCV, DA to membrane distances at the CCV gap were not considered in docking distance determinations. Membrane structures were classified as docked DAVs if the structure was present in at least two consecutive 3D planes, larger than 30 nm in diameter, and 30 nm or less from the DA distal end. DA-membrane docking analysis was independently performed by 3-5 different individuals for validation. The surface area of membrane structures docked to the MC was calculated using a built-in function Dragonfly tool.

### Immunoprecipitation

Immunoprecipitation was carried out as described^60^. 293T cells transfected with the specified plasmids were harvested and lysed using a lysis buffer (20□mM Tris-HCl, pH 7.5, 150□mM KCl, 1□mM EDTA, 0.5% NP-40, 10% glycerol, 10□mM sodium pyrophosphate, 3□mM dithiothreitol, and 0.5□mM phenylmethyl sulphonyl fluoride (PMSF, ThermoFisher, 36978) and protease inhibitor cocktail (Sigma, 539134). The resulting cell lysates were clarified through centrifugation at 14,000□g for 10□min at 4°C. Immunoprecipitation of GFP fusion proteins was performed using GFP-Trap affinity resin (Chromotek, gta-20), while HA-tagged proteins were immunoprecipitated with Pierce Anti-HA magnetic beads (ThermoFisher, 88836).Following four washes with the lysis buffer, the beads were incubated with 65□μl of 2× SDS loading buffer for subsequent western blot analysis.

For endogenous immunoprecipitation, 293T cells were grown to confluency in 10-cm plates. Following a 3 h serum starvation, the cells were lysed using a buffer composed of 50□mM Tris-HCl (pH 7.4), 150□mM NaCl, 1% Triton-X100, 10% glycerol, and a protease inhibitor cocktail. The cleared cell lysate was then mixed with 3 μg of EHD1 antibody or Rabbit-IgG, and the mixture was incubated for 1 h in a cold room. Subsequently, 20 μl of Affi-Prep Protein A Resin (BioRad) was added, and the samples were incubated for an additional 1 h. The beads were washed four times with lysis buffer and subjected to SDS-PAGE gel separation.

### Mass spectrometry (MS)

Before immunoprecipitation, cells were serum-starved for 3 h and then lysed with a lysis buffer containing 50□mM Tris-HCl (pH 7.4), 150□mM NaCl, 1% Triton-X100, 10% glycerol, and a protease inhibitor cocktail. The cleared cell lysates were mixed with 25 μl of GFP-Trap affinity resin (Chromotek, gta-20) and incubated for 2 h in a cold room. The resin was subsequently washed four times with the lysis buffer and then mixed with 50 μl of 50 mM NH4HCO3 for MS analysis.

### Statistics and reproducibility

Statistical analyses were performed using GraphPad Prism for Macintosh OS. Data presented are as specified in the Figure legends or text but generally ±SEM or SD. Two-or more group comparisons were carried out using an unpaired, two-tailed Student t-test or with One-way ANOVA as indicated Figure legends with significant values as follows: *P□<□0.05, **P□<□0.01, n.s. not significant, N number of independent experiments.

## Data availability

FIB-SEM datasets will be uploaded to EMPIAR (www.empiar.ebi.ac.uk) upon publication. Source data are provided within this paper.

## Acknowledgements

The authors thank Dr. Christine Insinna for critical reading of this manuscript. This project has been funded in part with Federal funds from the National Cancer Institute, National Institutes of Health, under Contract No. HHSN261200800001E. The content of this publication does not necessarily reflect the views or policies of the Department of Health and Human Services, nor does mention of trade names, commercial products, or organizations imply endorsement by the U.S. Government.

## Author Contributions

C.J.W. and Q.L. designed the study and wrote the manuscript. Q.L. conducted most of the cell biology and biochemistry experiments, with help from H.Z. on SRM and immunoprecipitation, and from V.M. and E.K. on ExM. Q.L. prepared the FIB-SEM samples; A.H. and K.N. collected the FIB-SEM images. A.S. generated vEM movies. C.J.W., Q.L., Z.K., A.S., K.N. and S.P. analyzed data.

## Ethics declarations

None to declare.

## Competing interests

The authors declare no competing interests.

## Supplementary information

Supplementary Figure 1-10 and Supplementary Table 1

Supplementary Movie 1. FIB-SEM and segmentation of primary cilium in RPE1 cells.

Supplementary Movie 2. FIB-SEM and segmentation of CV structure in human fibroblast cell.

Supplementary Movie 3. FIB-SEM and segmentation of CCV from a RPE1 cell.

Supplementary Movie 4. 3D analysis of CCV docking to the DA.

Supplementary Movie 5. 3D analysis of DAV docking to the DA. Supplementary Movie 6. Volume view of DAV-EMC structures in a RPE1 cell.

Supplementary Movie 7. 3D view of CCV and DAV structures on the MC of a RPE1 cell.

Supplementary Movie 8. SRM SIM^2^ live-cell imaging showing ciliogenesis progression.

Supplementary Movie 9. 3D view of TCV structure on the MC of a RPE1 cell.

## Notes

### Competing Interest Statement

The authors have declared no competing interest.

