## Supplementary File for "Characterization of membrane structures regulating primary ciliogenesis by quantitative isotropic ultrastructure imaging"

### Supplementary information

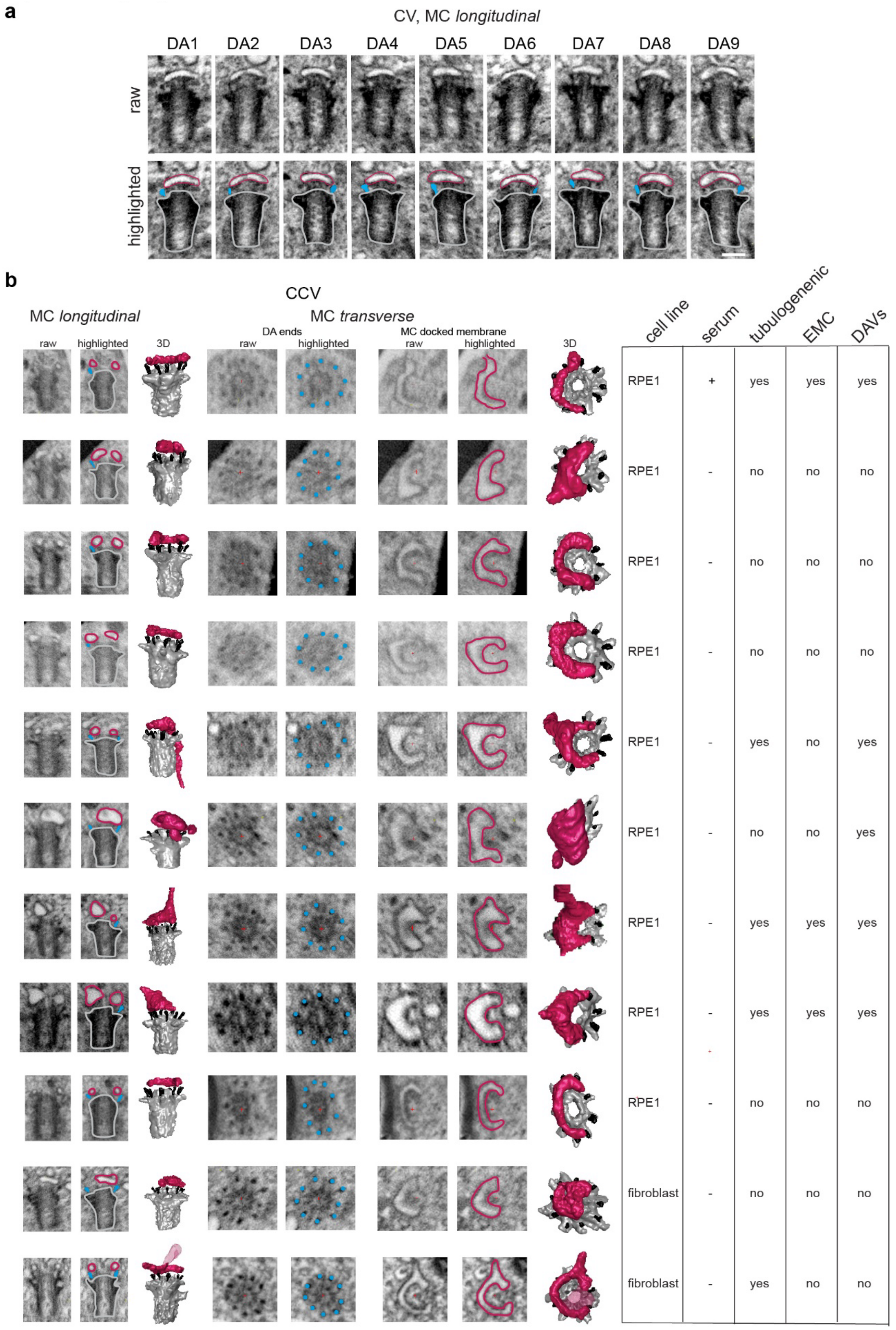


**Supplementary Fig. 1: Cilia and CCV structure reconstruction from FIB-SEM imaging.**

**Supplementary Fig. 1: Cilia and CCV structure reconstruction from FIB-SEM imaging.** **a** Longitudinal MC FIB-SEM images showing all 9 DAs and CV membrane shown in Fig. 1**c**. **b** CCV membrane structures identified at the MC in RPE1 and human fibroblast cells grown in serum or starved for 3 or 6h described in Fig. 1**b**. Transverse vEM images show the CCV structure directly above the DA ends and the longitudinal images show sections through the CCV appearing as DAV or CV structures. The presence of membrane tubule extensions and EMC connections to the PM are indicated for each cell.

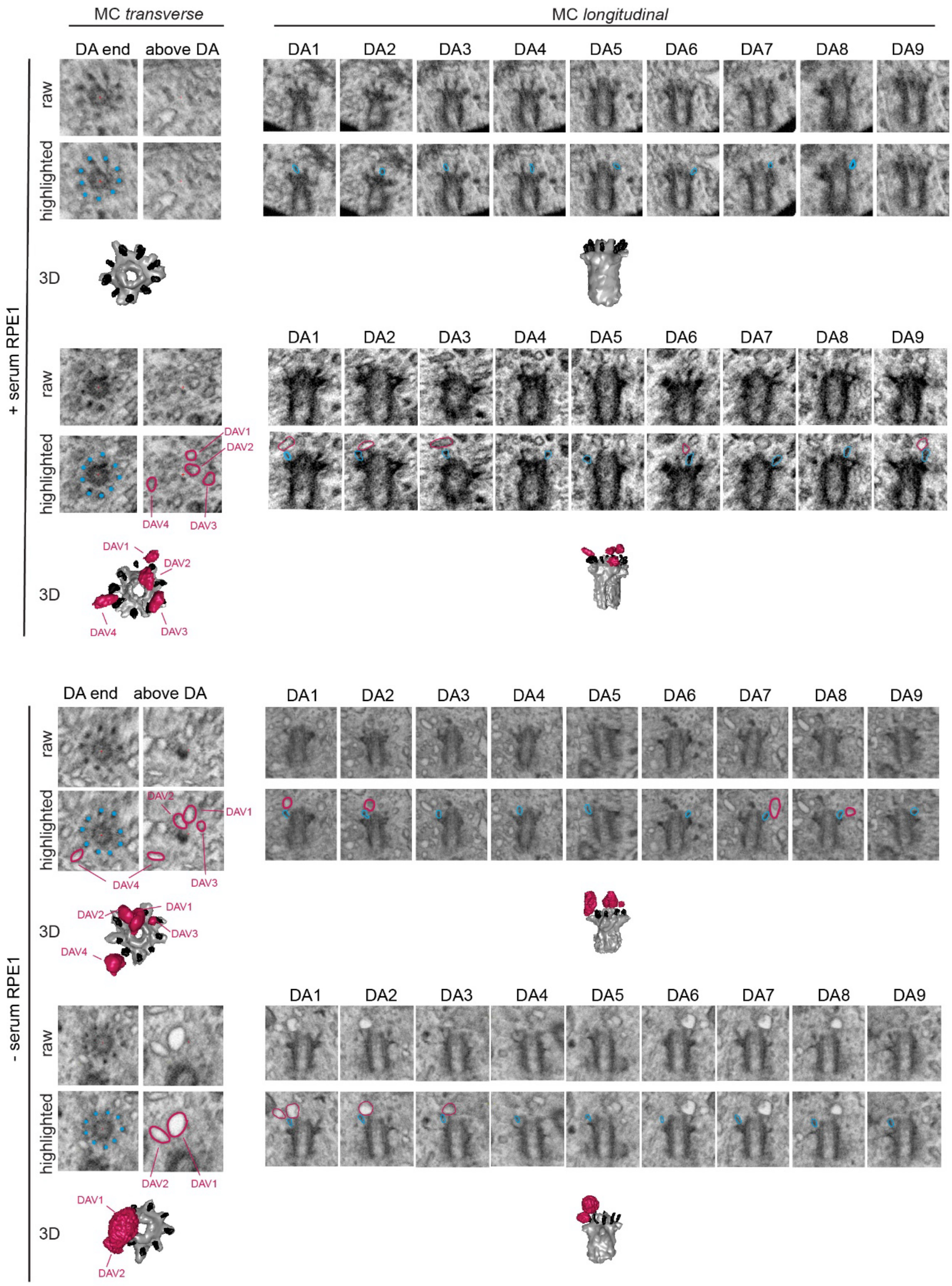


**Supplementary Fig. 2: Determination of DAV docking to the MC.**

**Supplementary Fig. 2: Determination of DAV docking to the MC.**

FIB-SEM and segmented images mages showing DAs and docked DAVs in serum-fed and serum-starvation conditions for RPE1 cells described in Fig. 2**d**. Longitudinal FIB-SEM images show each of the 9 DAs and association DAVs and transverse sections show the MC DA ends and associated DA-membrane docking. Highlighted structures DAV (magenta), DAs (cyan) and MC (grey) shown in the unmarked raw image.

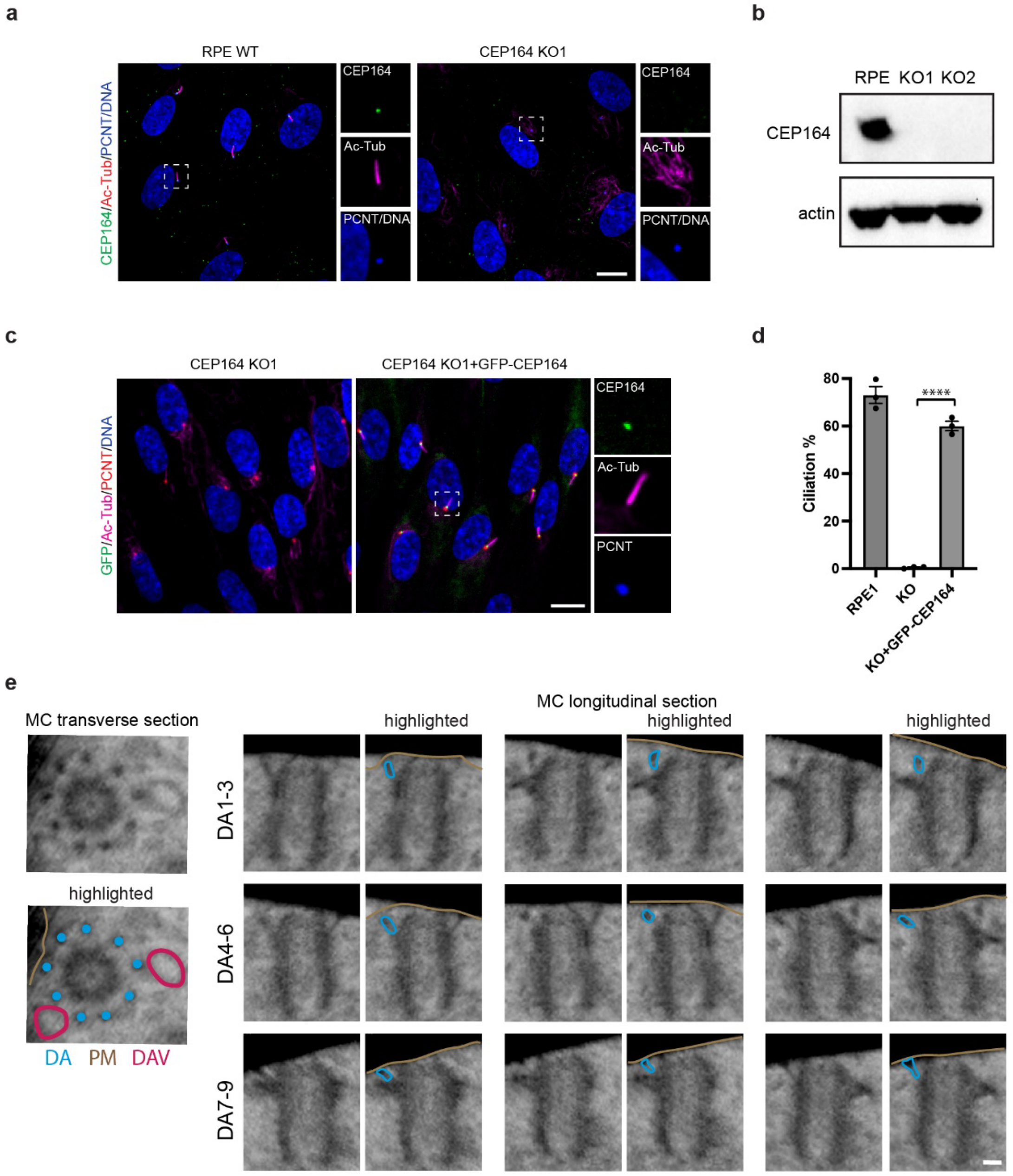


**Supplementary Fig. 3: CEP164 KO ciliogenesis analysis in RPE1 cells.**

**Supplementary Fig. 3: CEP164 KO ciliogenesis analysis in RPE1 cells.**

**a** RPE1 WT and CEP164 KO cells were serum starved 24h and immunostained with CEP164, $^{Ac}$tub and pericentrin antibodies and the nuclear marker Hoechst. Scale bars 10 µm.

**b** Immunoblot of CEP164 protein expression in WT RPE1 and CEP164 KO cells using CEP164 and actin antibodies.

**c**, **d** GFP-CEP164 expression rescues ciliogenesis in CEP164 KO RPE1 cells. Cells were treated as described in **a** and cilia levels quantified. Means ± SEM (3 independent experiments, n: RPE1=462 cells, KO=380 cells, KO+GFP-CEP164=583 cells), ****P<0.0001, Scale bars 10 µm.

**e** Partial docking of MC to the PM in CEP164 KO cells. FIB-SEM images of CEP164 depleted RPE1 GFP-CETN1 cells serum starved for 24h showing partial PM docking shown in Fig. 3a. DA 3 and 4 are not docked to the PM and DAVs are detected. Traced DAs (cyan), PM (brown) and DAVs (magenta) for corresponding FIB-SEM sections are shown. Scale bar 100 nm.

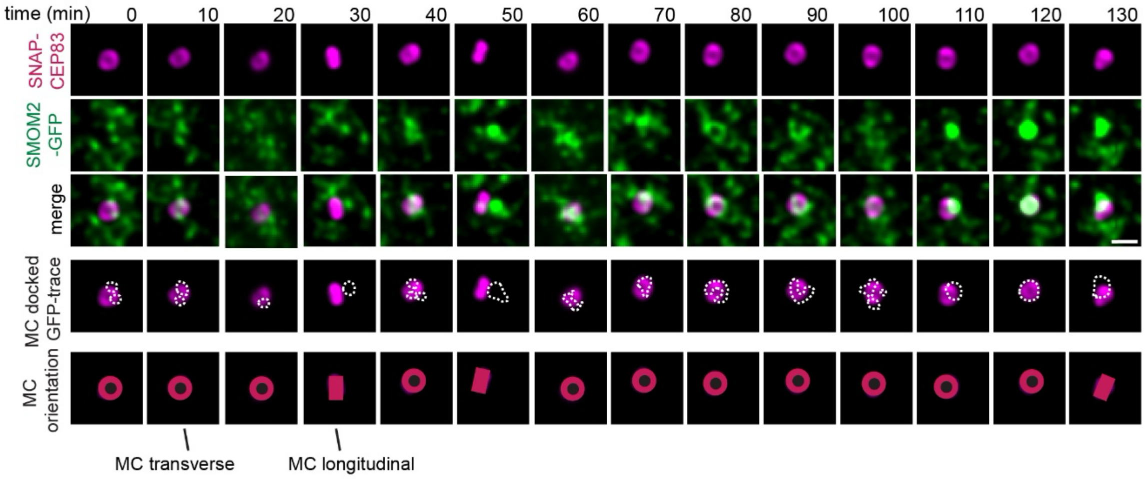


**Supplementary Fig. 4: SRM live cell imaging of ciliogenesis membrane assembly.**

SIM$^2$ super-resolution live cell imaging of RPE1 cell as described in Fig. 4d. Scale bar 1 µm. Bottom panels show the positional orientation of the MC DAs (magenta illustration) in the imaging plane. White doted lines indicate the outline of membrane vesicles. Ring = MC facing up, solid line = MC sideways.

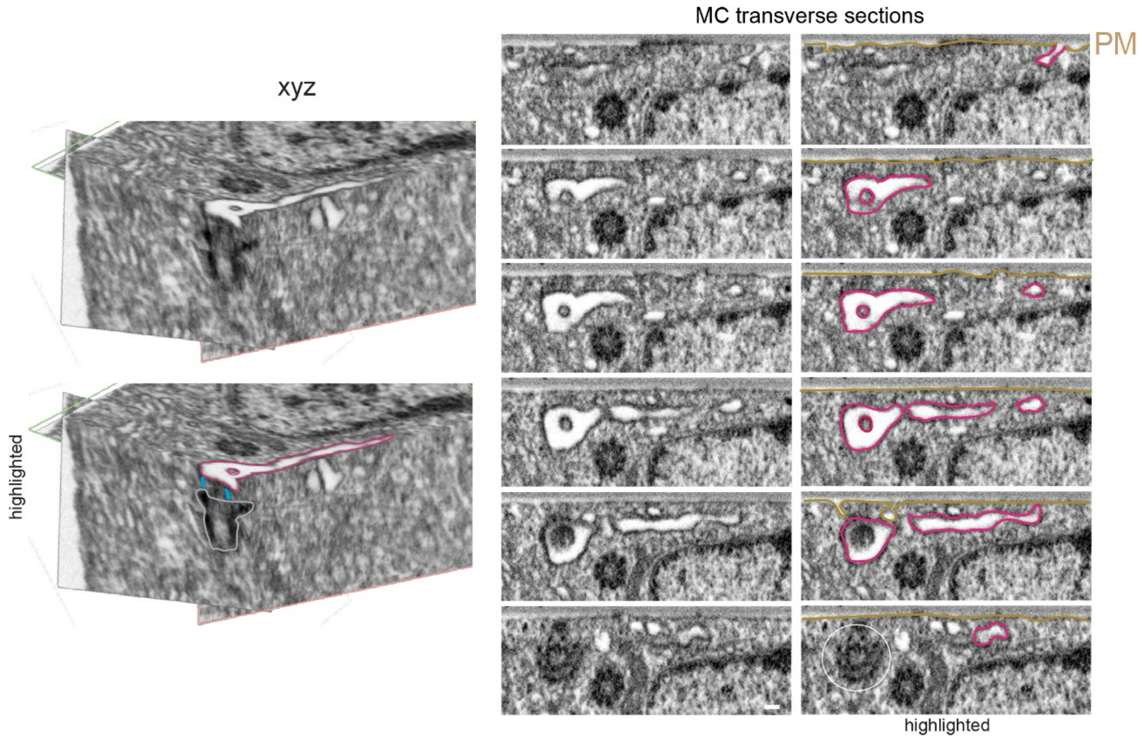


**Supplementary Fig. 5: FIB-SEM image slices of TCV.**

FIB-SEM images for the toroidal membrane shown and described in Fig. 4e. (right images) Images show xyz planes through MC, toroid and EMC. (left image) additional transverse sections through MC and enlarged area showing the EMC connection to the PM. White circle shows MC position. Scale bar = 100 nm.

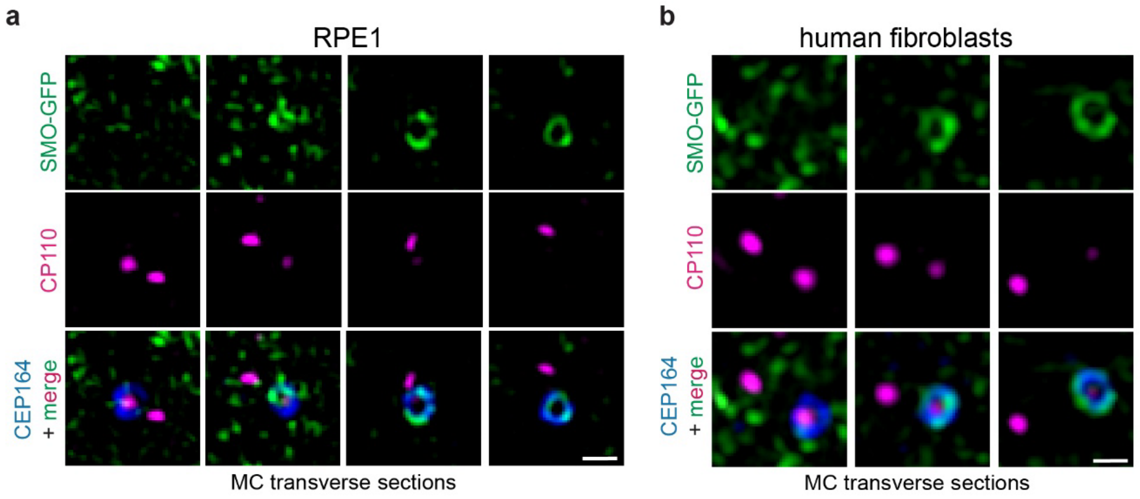


**Supplementary Fig. 6: MC uncapping association with ciliogenesis membrane accumulation.**

**a**, **b** Accumulation of SMO-GFP and CP110 on the MC in 6h serum starved RPE1 (**a**) and human fibroblast (**b**) cells. Cells stably expressing SMO-GFP cells were fixed and stained with CP110 and CEP164 antibodies and imaged by Nikon SIM. Scale bars 500 nm.

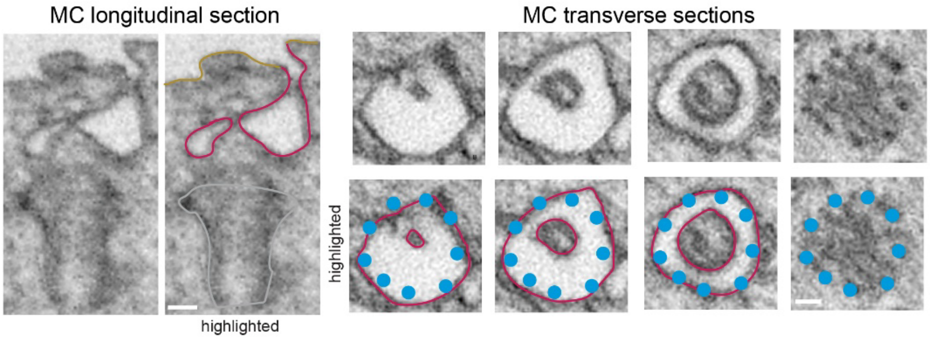


**Supplementary Fig. 7: FIB-SEM image slices of TCV.**

Transverse (right 4 images) and longitudinal (left 2 images) FIB-SEM images for the cell shown in the bottom panel of Fig. 5a with CP110 removed from the. MC showing a TCV docked to the MC  Transverse sections shows the presence of an EMC. PM highlighted in gold, TCV and/or EMC are highlighted in magenta and DA ends are marked with a cyan dot. Scale bars = 100 nm.

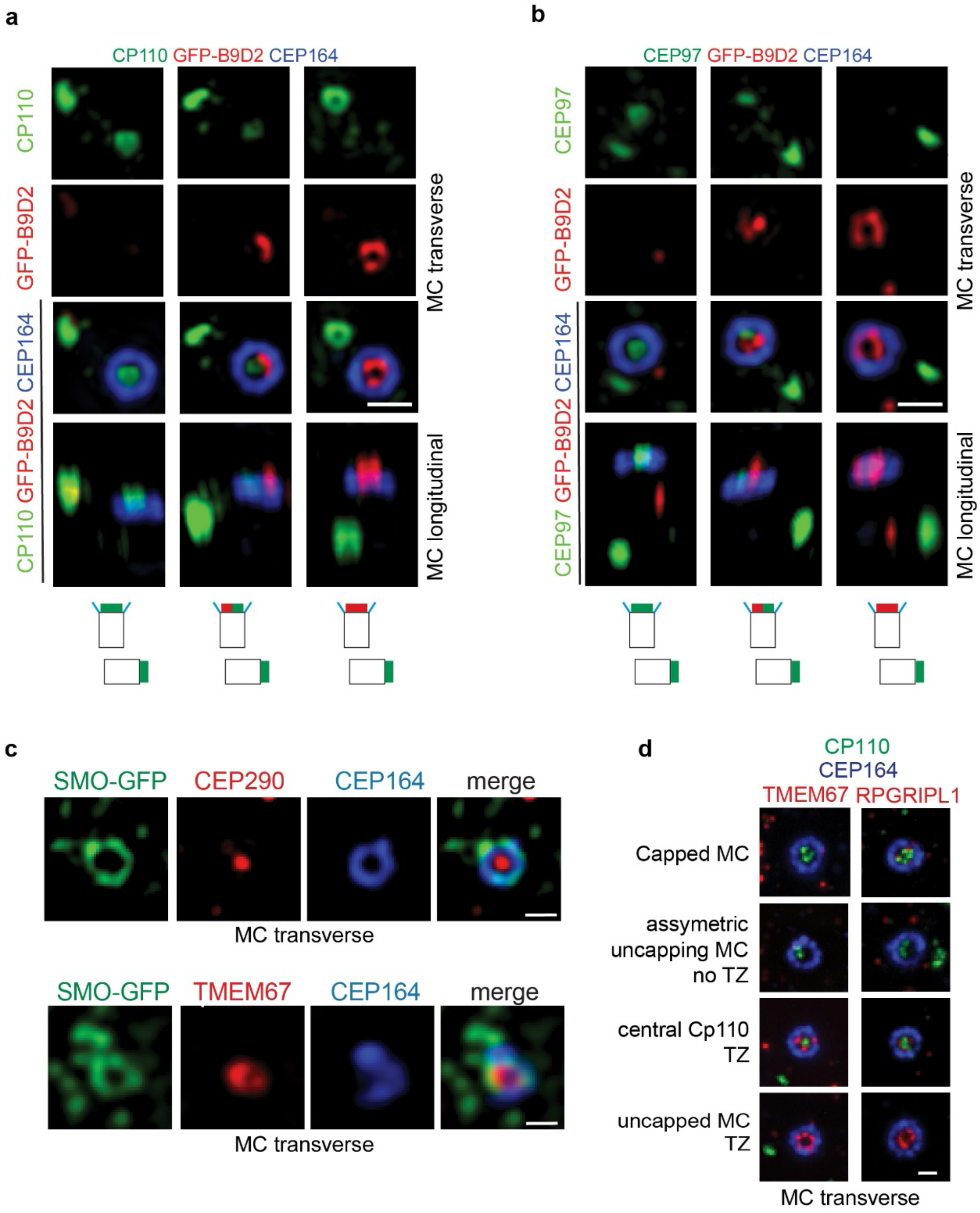


**Supplementary Fig. 8: MC uncapping association with TZ protein recruitment.**

**a**, **b** 3D-SIM images showing transition zone protein B9D2 recruitment to the MC in relation to MC uncapping during ciliogenesis. RPE1 cells stably expressing GFP-B9D2 were serum starved for 6h and stained with CEP164 and MC cap proteins CP110 (**a**) and CEP97 (**b**). Cartoons show fluorescence marker position on MC and DC associated with CP110 removal and TZ protein recruitment. Cells were imaged by OMX-SIM. Scale bars = 500 nm.

**c** SIM images showing TZ protein MC localization in RPE1 cells with toroidal-like membranes. Cells expressing SMO-GFP were serum starved 6h and stained with TMEM67 or CEP290. SMO-GFP ring structures at the distal end of the MC. Scale bars = 500 nm.

**d** Representative images showing observed CP110 and TZ protein localization at the MC described in plots for Fig. 5d, e. Scale bar = 1 μm.

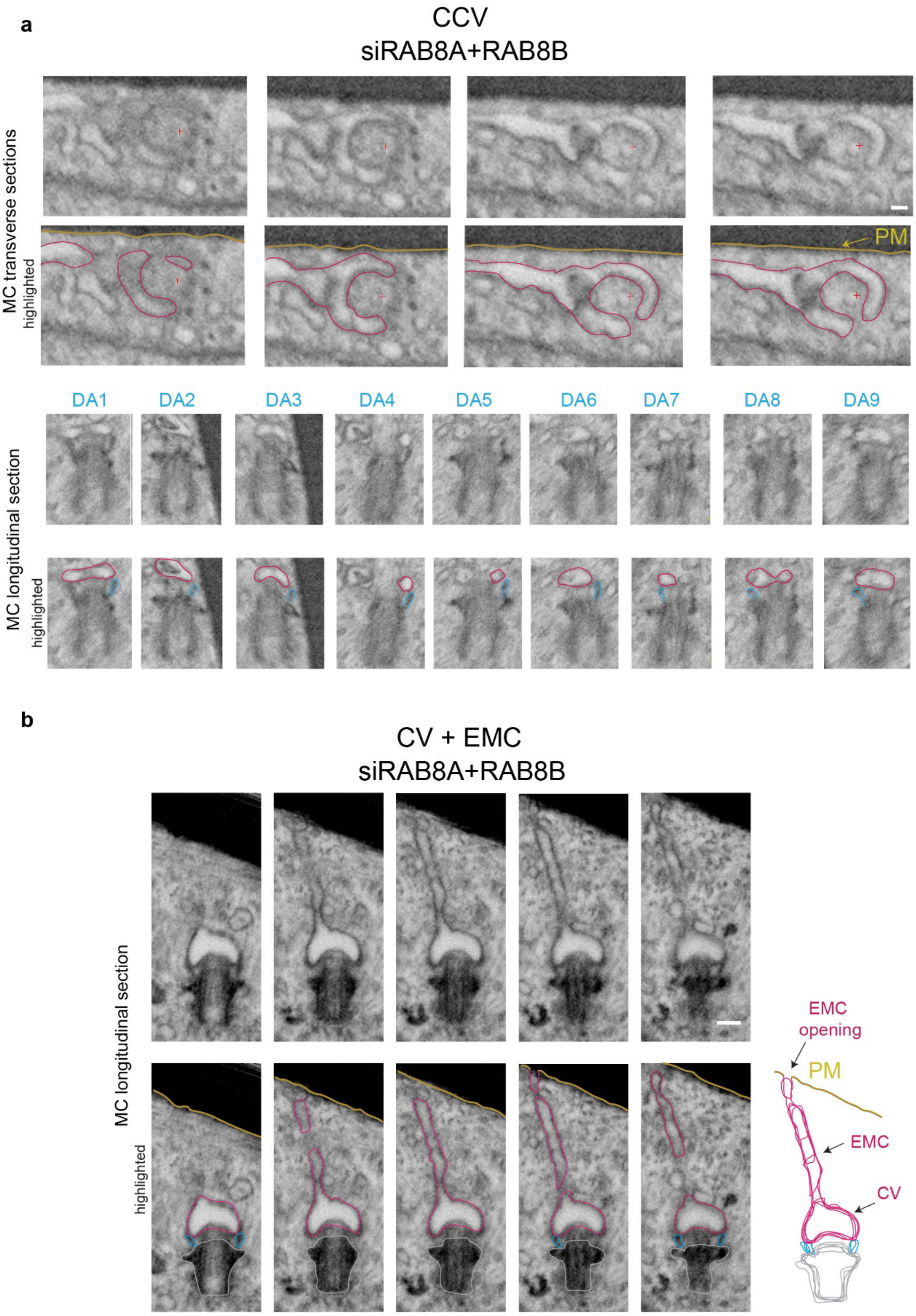


**Supplementary Fig. 9: FIB-SEM images of RAB8 depleted cell showing a CCV.**

**Supplementary Fig. 9: FIB-SEM images of RAB8 depleted cell showing a CCV.**

**a** Transverse (top panels) and longitudinal (bottom panels) MC FIB-SEM images shown for cell described in Fig. 7c with CCV and extended membrane tubule. All 9 DA are shown in longitudinal sections (cyan trace) and CCV-membrane tubule (magenta). Plus (+) is a positional marker for section. Scale bar = 100 nm.

**b** MC longitudinal FIB-SEM images shown for cell shown in Fig. 7e showing CV-EMC (magenta) connection to PM (gold trace). Scale bar = 200 nm.

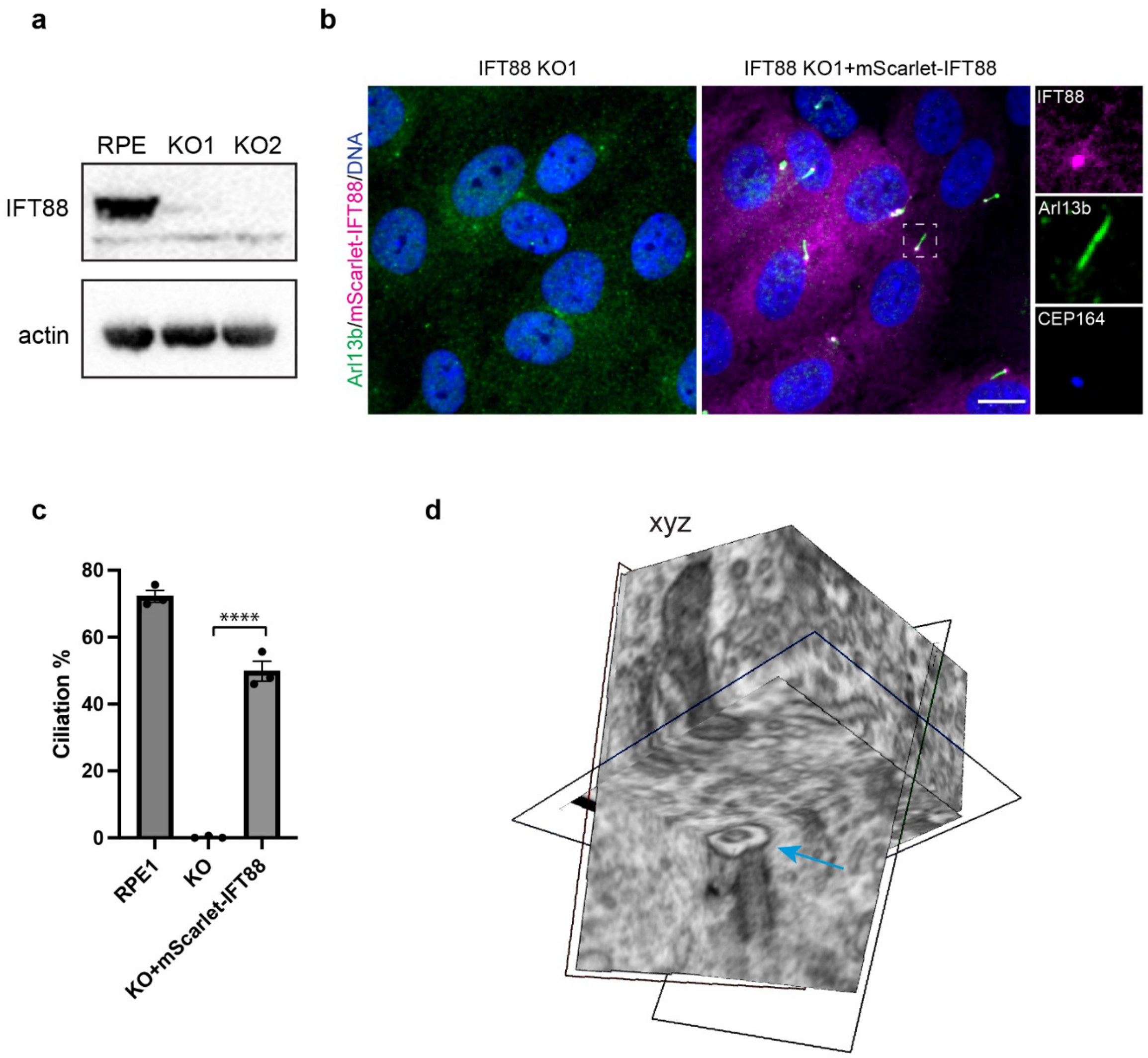


Supplementary **Fig. 10: RPE1 cell IFT88 KO ciliation analysis.**

**a** Immunoblot of IFT88 protein expression in wild type RPE1 and IFT88 KO cells using IFT88 and actin antibodies.

**b**, **c** mScarlet-IFT88 expression rescues ciliogenesis in IFT88 KO RPE1 cells. Cells were serum starved 24h and immunostained with Arl13b (cilia) and CEP164 antibodies and ciliation quantified. Means ± SEM (3 independent experiments, n: RPE1=521 cells, KO=515 cells, KO+mScarlet-IFT88=490 cells). ****P<0.0001. Scale bar 10 μm.

**d** FIB-SEM x,y,z planes of membrane toroid shown in Fig. 8d. Blue arrow shows MC structure.

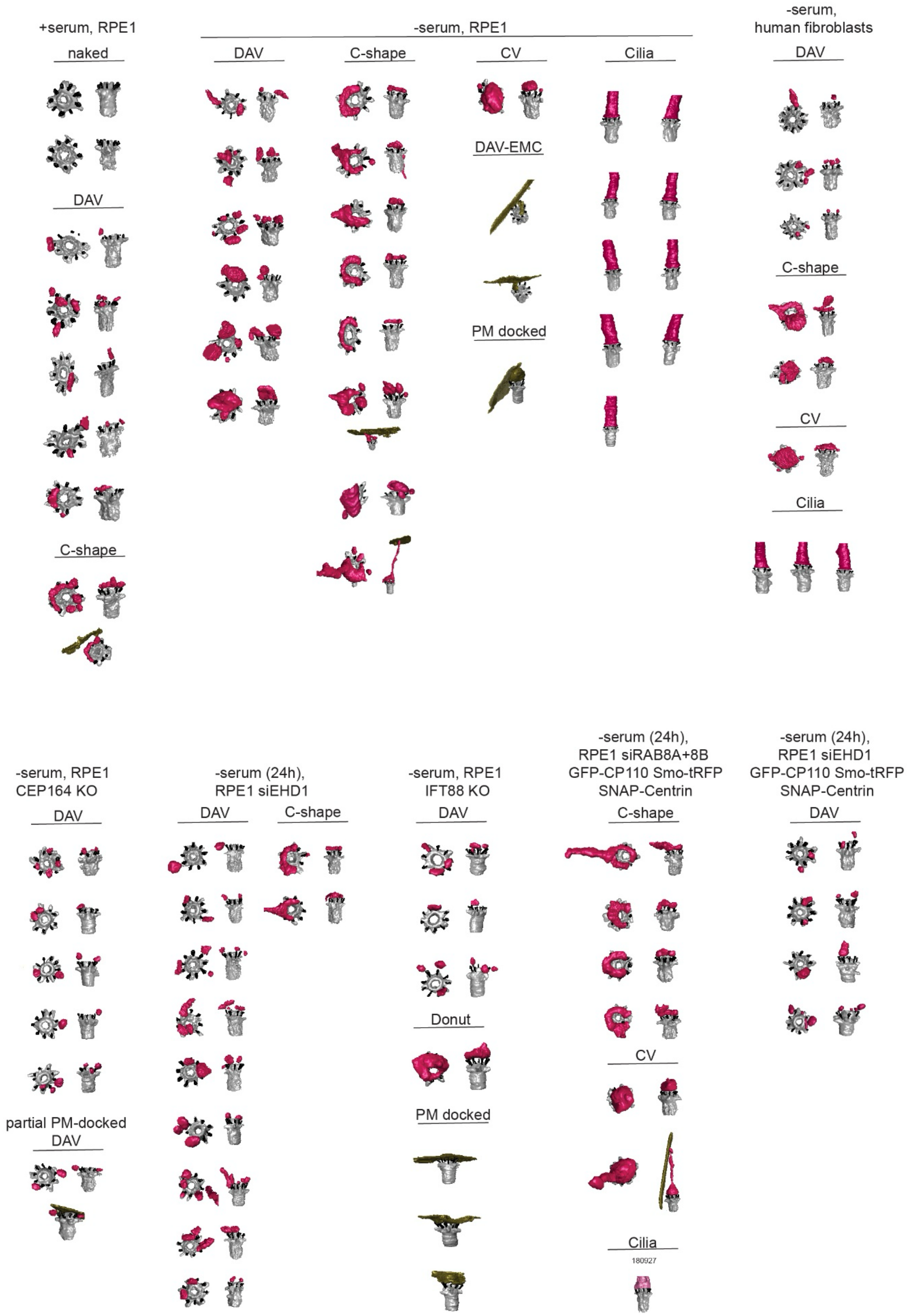


**Supplementary Table 1. FIB-SEM segmented MC structures.**

**Supplementary Movie 1. FIB-SEM and segmentation of primary cilium in RPE1 cells.**

3D longitudinal FIB-SEM and segmentation images of a RPE1 cilium shown in Fig. 1a. DA are highlighted in cyan and ciliary membrane in magenta.

**Supplementary Movie 2. FIB-SEM and segmentation of CV structure in human fibroblast cell.**

3D FIB-SEM and segmentation images of CV for the human fibroblast cell shown in Fig. 1c.

**Supplementary Movie 3. FIB-SEM and segmentation of CCV from a RPE1 cell.**

3D FIB-SEM and segmentation images of CCV from a RPE1 cell**.**

**Supplementary Movie 4. 3D analysis comparing cilia and CCV membrane docking to the DA.**

Segmented cilia from Supplementary Movie 1 (left) and CCV Supplementary Movie 3 (right) showing 3D analysis of membrane docking at the DA. 3D blue mesh spheres show 30 nm radius from the DA ends.

**Supplementary Movie 5. 3D analysis of DAV docking to the DA.**

3D FIB-SEM and segmentation showing analysis of DAV (magenta) and non-docked membranes (yellow). Analyzed DA are highlighted from black to cyan coloration and DAVs are identified within 30 nm of the DA ends (blue mesh sphere).

**Supplementary Movie 6. Volume view of DAV-EMC structures in a RPE1 cell.**

3D FIB-SEM and segmentation images of DAV-EMC containing MC shown in Fig. 2e. DAV-EMC structure is highlighted from gold to magenta coloration. PM associated membranes are colored in gold.

**Supplementary Movie 7. 3D view of CCV and DAV structures on the MC of a RPE1 cell.**

3D FIB-SEM and segmentation images of a MC containing a CCV and DAVs from RPE1 cells**.** Blue mesh spheres show the 30 nm radius from the end of the DA.

**Supplementary Movie 8. SRM SIM$^2$ live-cell imaging showing ciliogenesis progression.**

SRM SIM$^2$ live cell time-lapse imaging showing ciliogenesis progression as in Fig. 4d. RPE1 cells expressing SMOM2-GFP and SNAPf-CEP83 were stained with SNAP-Cell 647-SiR dye and imaged every 10 min after serum-starvation. Images showing max-intensity projection of z-stacks around the basal body. Scale bar 500 nm.

**Supplementary Movie 9. 3D view of TCV structure on the MC of a RPE1 cell.**

3D longitudinal FIB-SEM and segmentation images of the TCV shown in Fig. 4e. DA are highlighted in cyan and TCV in magenta.
